# Improved control of Septoria tritici blotch in durum wheat using cultivar mixtures

**DOI:** 10.1101/664078

**Authors:** S. Ben M’Barek, P. Karisto, W. Abdedayem, M. Laribi, M. Fakhfakh, H. Kouki, A. Mikaberidze, A. Yahyaoui

## Abstract

Mixtures of cultivars with contrasting levels of resistance can suppress infectious diseases in wheat, as demonstrated in numerous field experiments. Most studies focused on airborne pathogens in bread wheat, while splash-dispersed pathogens have received less attention, and no studies have been conducted in durum wheat. We conducted a two-year field experiment in Tunisia, to evaluate the performance of cultivar mixtures with varying proportions of resistance (0–100%) in controlling the polycyclic, splash-dispersed disease Septoria tritici blotch (STB) in durum wheat. To measure STB severity, we used a high-throughput method based on digital image analysis of 3074 infected leaves collected from 42 and 40 experimental plots during the first and second years, respectively. This allowed us to quantify pathogen reproduction on wheat leaves and to acquire a large dataset that exceeds previous studies with respect to accuracy and precision. Our analyses show that introducing only 25% of a disease-resistant cultivar into a pure stand of a susceptible cultivar provides a substantial reduction of almost 50% in disease severity compared to the susceptible pure stand. However, comprising the resistant component of two cultivars instead of one did not further improve disease control, contrary to predictions of epidemiological theory. Susceptible cultivars can be agronomically superior to resistant cultivars or be better accepted by growers for other reasons. Hence, if mixtures with only a moderate proportion of the resistant cultivar provide a similar degree of disease control as resistant pure stands, as our analysis indicates, such mixtures are more likely to be accepted by growers.

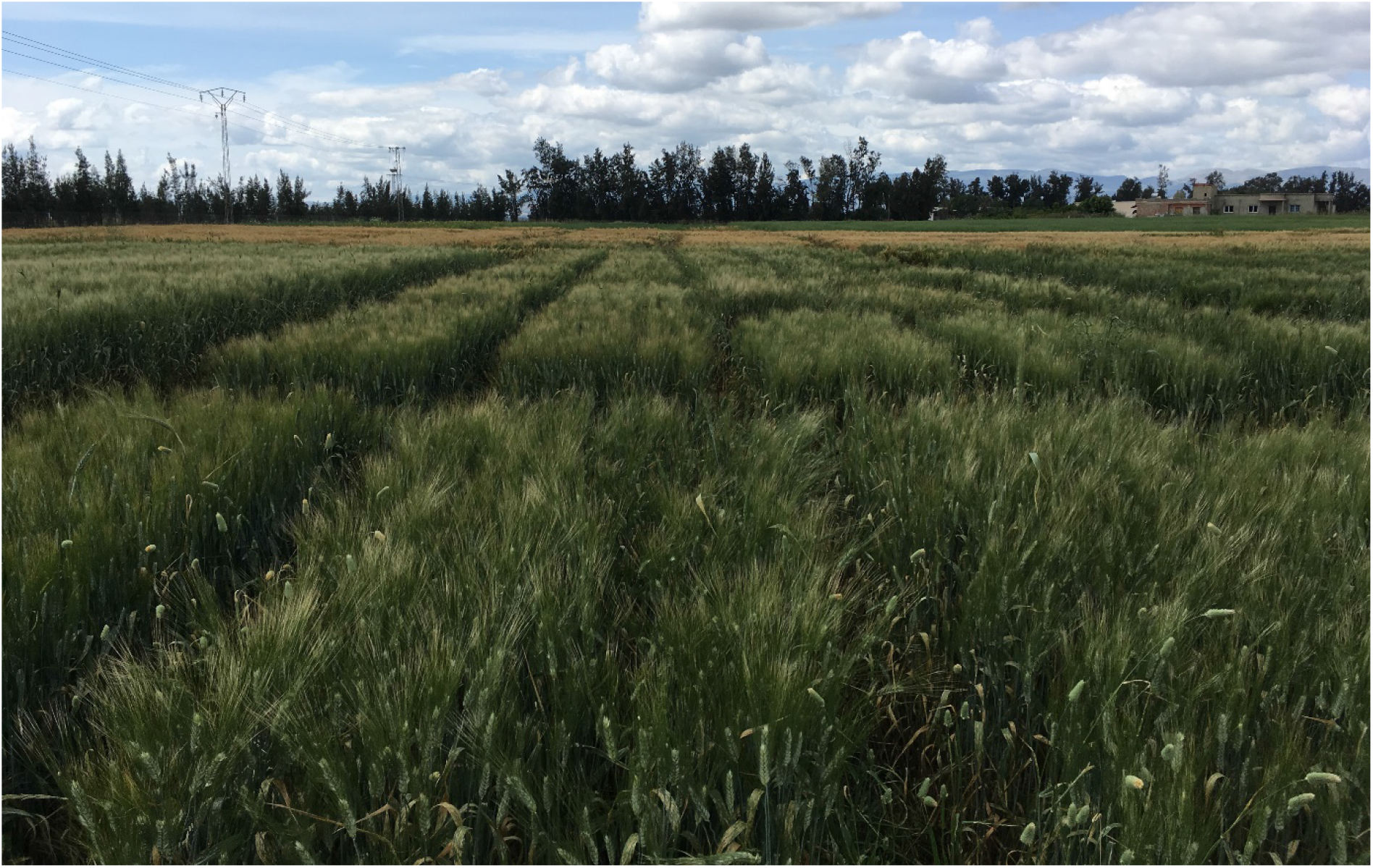

## Introduction

A deliberate introduction of genetic diversity into crop plant populations has been proposed as a promising way to suppress plant disease epidemics (Wolfe, 1985; Finckh et al., 2000; Mundt, 2002) and enhance the overall crop function (Newton et al., 2009). One way to diversify crop plants is to grow two or more genetically distinct cultivars of the same crop concurrently within the same field. This can be achieved by mixing seeds of different cultivars before sowing, thereby creating a physical cultivar mixture.

The idea behind cultivar mixtures is that genetic, physiological, structural, and phenological diversity among the components of the mixture (i.e., among different cultivars that comprise the mixture) may drive beneficial interactions not only between cultivars but also between cultivars and environments (Kiær et al., 2009; Newton et al., 2009; Borg et al., 2018). As a result, cultivar mixtures have proven to improve the resilience to biotic and abiotic stresses in crops and boost yield and its stability compared to pure stands, especially in low-pesticide cropping systems (Kiær et al., 2009; Smithson and Lenné, 1996; Borg et al., 2018). Mixtures can also enhance product quality, if the components of the mixture are chosen appropriately (Finckh et al., 2000; Mundt, 2002). For these reasons, cultivation of cultivar mixtures has gained interest in several countries (Borg et al., 2018; de Vallavieille-Pope et al., 2006; Finckh and Wolfe, 1997; Wolfe et al., 2008).

Cultivar mixtures suppress the development of disease epidemics when the mixture components have contrasting levels of resistance to the targeted disease (Wolfe, 1985; Finckh and Wolfe, 2006; Gigot et al., 2013). Consequently, most studies investigated mixtures of disease-susceptible and disease-resistant cultivars (cf. Garrett and Mundt, 1999, and Mikaberidze et al., 2015, for discussion of mixtures that contain two or more resistant cultivars). The most important mechanisms of disease reduction in cultivar mixtures are the dilution (or density) effect, the barrier effect, induced resistance (Wolfe, 1985; Finckh et al., 2000), and competition among pathogen strains (Garrett and Mundt 1999). From an evolutionary perspective, appropriately designed mixtures are expected to hamper adaptation of the pathogen and increase the durability of the resistance genes deployed (Finckh et al., 2000; Mundt, 2002).

Many studies presented convincing empirical evidence that cultivar mixtures provide effective control of airborne cereal diseases, particularly rusts and mildews, as reviewed by Wolfe (1985), Finckh et al. (2000), and Mundt (2002). See also a meta-analysis of 11 publications on stripe (yellow) rust of wheat (Huang et al., 2012). However, effect of cultivar mixtures on splash dispersed cereal diseases is less studied and the disease reduction by mixtures appears less consistent and is on average lower in magnitude compared to airborne diseases (Jeger et al., 1981b; Mundt et al., 1994; Newton et al., 1997; Garrett and Mundt, 1999).

Here, we investigated Septoria tritici blotch (STB) disease, which is predominantly splash dispersed. STB is one of the major threats to wheat production worldwide. It is caused by the fungal pathogen *Zymoseptoria tritici.* The infection cycle of the fungus starts when asexual pycnidiospores or airborne sexual ascospores land on a susceptible wheat leaf. The asymptomatic phase lasts for about 8-14 days. The switch to necrotrophy leads to a collapse and death of the host mesophyll cells usually between 12 and 25 days after penetration (Karisto et al., 2019a). Within necrotic lesions, the fungus begins to reproduce asexually and later sexually (Ponomarenko et al., 2011). Under conducive conditions, this polycyclic pathogen can complete up to six asexual infection cycles during one growing season. Because of *Z. tritici*’s mixed reproductive system, large population sizes and longdistance dispersal, its populations are extremely diverse (Linde et al., 2002; Hartmann et al., 2018).

Tunisia is a key durum wheat producer in the Mediterranean region but is also the largest per capita wheat consumer in the world (FAO, 2017). The main growing areas of durum wheat are located in the Northern part of the country under rain-fed conditions of a subhumid climate (Ammar et al., 2011), favourable for fungal diseases. Among these, STB poses an especially serious threat to Tunisian wheat production. Control of STB in Tunisia relies largely on fungicides and resistant cultivars. The use of chemical compounds has been adopted by Tunisian durum wheat growers at a slower pace as compared to bread wheat growers in Europe. At the same time, chemical control is costly and poses substantial risks to the environment and human health. Both factors give the farmers an incentive to reduce or avoid the use of fungicides.

The vast majority of commercial cultivars in Tunisia are highly susceptible to STB and new, STB-resistant cultivars are released at a very slow rate (Ammar et al., 2011). The variety Karim, released in 1980, covers more than 60% of the durum wheat acreage (Ammar et al., 2011; Rezgui et al., 2008). Even though highly susceptible to STB, favourable agricultural properties combined with its relatively low gluten and low yellow flour content (Ammar et al., 2011) made it the farmer’s favourite. During the last decade, a few STB-resistant cultivars were released in Tunisia, including ‘Salim’ that was registered in 2010. At the time of release, Salim showed resistance to STB (Gharbi and El Felah, 2013), but it is gradually becoming more susceptible (Bel Hadj Chedli et al., 2018). In 2017, another promising variety, INRAT100, that contained Salim in its pedigree, was shown to be productive and resistant to several diseases including Septoria and Powdery mildew (Hammami and Gharbi, 2018). Furthermore, among recently imported cultivars released in Tunisia, ‘Monastir’ is resistant to STB (SOSEM, 2018; Bel Hadj Chedli et al., 2018).

Currently, the two methods to control STB, fungicides and a few resistant cultivars, available to Tunisian farmers are not providing satisfactory levels of disease control. The country experiences serious recurrent epidemics of STB with yield losses reaching up to 40% (Berraies et al., 2014). A large part of the reason is that many farmers simply do not use either of the two control methods and neglect crop rotation practices. In addition, farmers often cannot afford fungicides. Those farmers who do use either of the two control methods exert a strong directional selection on pathogen populations. As a result, both fungicides and resistant cultivars are likely to rapidly lose efficacy against *Z. tritici* that has a high evolutionary potential (McDonald and Linde, 2002). Thus, there is an urgent need to re-consider the way STB is controlled in Tunisia and devise management strategies that are not only efficient and sustainable, but also likely to be accepted by growers. Cultivar mixtures may prove to be such a strategy that is particularly well-suited to Tunisian conditions.

Despite the importance of durum wheat, the bulk of research on host resistance to STB has been conducted in bread wheat *(Triticum aestivum)* (Kollers et al., 2013; Brown et al., 2015; Karisto et al., 2018; Yates et al., 2019). At present, our understanding of the genetics and molecular basis of STB resistance in durum wheat is limited. However, this situation may improve with the recent studies on Ethiopian durum wheat landraces (Kidane et al., 2017), Tunisian landraces (Aouini, 2018), and the publication of the fully assembled durum wheat genome (Maccaferri et al., 2019).

Similarly, cultivar mixtures for controlling STB have been studied in bread wheat but not in durum wheat (Mundt et al., 1995; Mille and Jouan, 1997; Cowger and Mundt, 2002; Mille et al., 2006; Gigot et al., 2013; Vidal et al., 2017). In most cases, STB severity in cultivar mixtures is moderately reduced compared to the expectation based on the severity in the pure stands, although Cowger and Mundt (2002) did not observe a consistent pattern. In particular, Gigot et al. (2013) found that a susceptible cultivar was consistently protected in a mixture under low to moderate STB levels. Subsequent greenhouse experimentation (Vidal et al., 2017) and a modelling study (Vidal et al., 2018) have demonstrated the importance of canopy structure for the efficacy of a mixture.

The previous studies have not systematically investigated the effect of the proportion of different cultivars in the mixture (mixing proportion) on its efficacy against STB. Investigation of mixing proportions is important both fundamentally and practically. Fundamentally, it provides a more robust and reliable measure of mixture efficacy across the whole range of mixing ratios, which may help to distinguish between different mechanisms of disease reduction in mixtures. From a practical perspective, growers may accept more easily mixtures that contain a small proportion of the resistant cultivar but still provide satisfactory disease control.

The number of different cultivars, or components, that comprise a mixture is another important parameter that affects the efficacy of disease control. Mille and Jouan (1997) and Mille et al. (2006) considered mixtures with more than two components to control STB, but they only used equal proportions of cultivars in mixtures. For this reason, the proportion of the susceptible component varied between two-way, three-way, and four-way mixtures. This precluded consistent testing of whether more than one resistant component in a mixture further improves disease control, as predicted by epidemiological modelling (Mikaberidze et al., 2015).

The objectives of this study were to fill the gaps in current knowledge identified above. We investigate how the proportion of the cultivars in the mixture influences the efficacy of the mixture in controlling STB. The aim is to find the lowest effective proportion of resistance. Additionally, we determine whether a mixture that contains two resistant components improves the control of STB compared to a mixture with only one resistant component.

## Materials and methods

### Study area and experimental design

Field experiments were conducted during the 2017–2018 and 2018–2019 wheat growing seasons at the CRP Wheat Septoria Precision Phenotyping Platform – experimental station of Kodia, which is located in the semi-arid region (36°32’51.89”N, 9°0’40.73”E, governorate of Jendouba, Tunisia, Figure 1d). The average annual rainfall in this area varies from 400 to 500 mm and the temperature usually ranges between 9.8°C (average minimum temperature) and 33°C (average maximum temperature). Daily temperature in the governorate of Jendouba and daily precipitation at the experimental station are shown in Figure S1 for both years of experiments. This region is considered to be a natural hot spot for STB disease (Bel Hadj Chedli et al., 2018).

**Figure 1.**
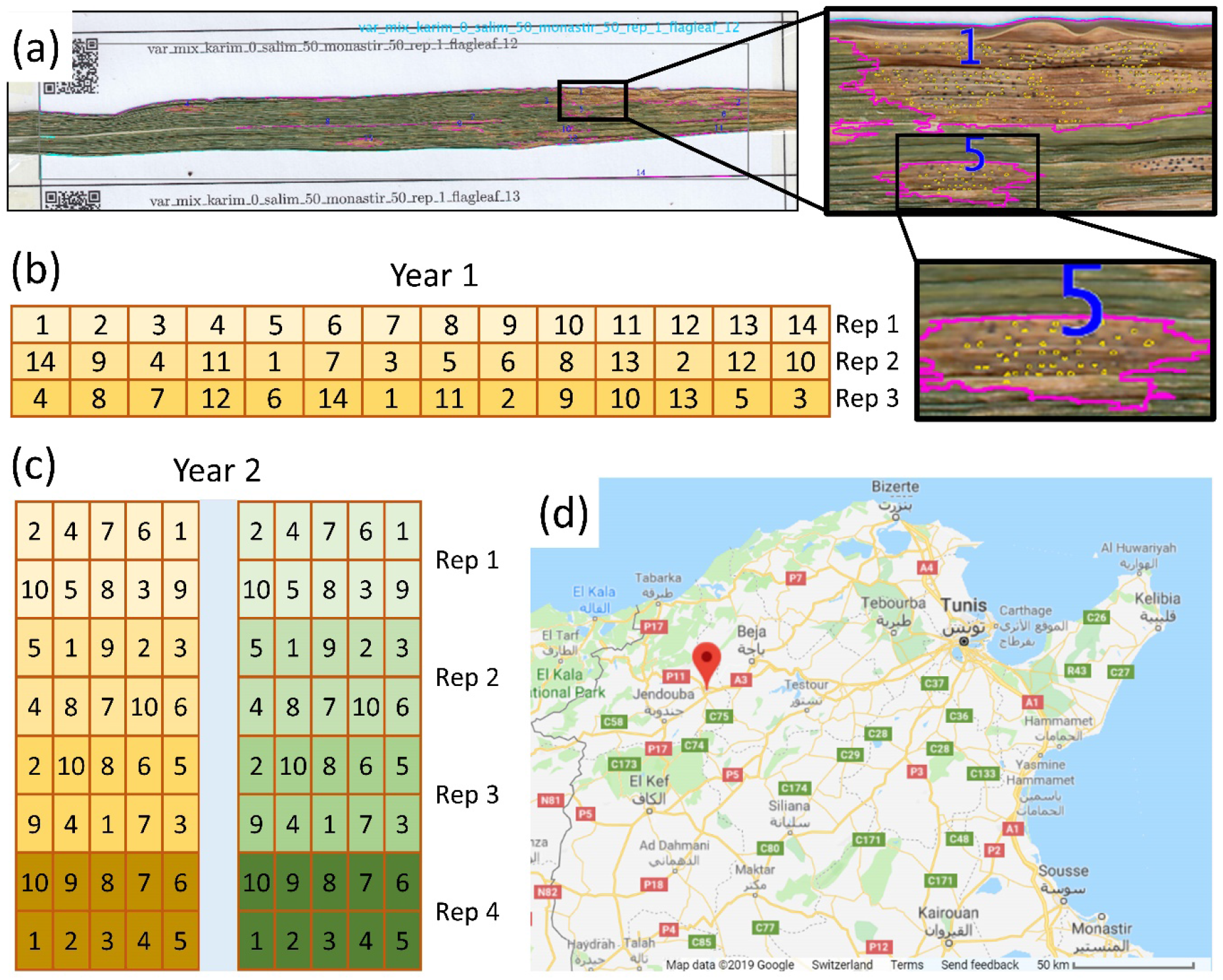
(a) An example of the automated leaf image analysis. Data is assigned to leaf labels based on the QR codes. Cyan lines represent the leaf boundary, purple lines show lesion boundaries, and yellow circles represent detected pycnidia. Very small lesions are not analysed. (b) Experimental field layout, the first year. Orange: experimental plots, numbers correspond to different treatments (Table 1), each of them replicated three times. (c) Experimental field layout, the second year. Orange: inoculated plots, green: uninoculated fungicide treated plots, numbers correspond to different treatments (Table 1), each of them replicated four times. (d) Location of the experimental site. The map from Google.

For the first year, three commercial durum wheat cultivars (Karim, Salim and Monastir) were chosen based on their contrasting scores of resistance to STB, similar earliness and plant height. The susceptible cultivar Karim was originally selected from an introduced F4 bulk from CIMMYT. Cultivar Salim was selected from a cross made in Tunisia in 1993. Cultivar Monastir was imported from France and released in 2012 (SOSEM, 2018). In addition to the three cultivars and their mixtures, we included pure stands of INRAT100, a promising variety that was registered in 2017 but is not yet released. There were 14 treatments in total: four pure stands, seven two-way mixtures and three three-way mixtures (Table 1). The mixtures with Karim contained always 25%, 50% or 75% of Karim, the rest being resistant cultivars. Each treatment was replicated three times in different plots.

**Table 1.** Proportions of cultivars in pure stands, two-way and three-way mixtures planted in three replicates in 2017/2018 cropping season (Year 1) and in 2018/2019 cropping season (Year 2) at the Septoria Phenotyping Platform -- Kodia station (Bou Salem, Tunisia). Treatment numbers correspond to numbers in Fig. 1.

| Year 1 |  | Pure stands |  |  |  | Two-way mixtures |  |  |  |  |  | Three-way |  |  |
| --- | --- | --- | --- | --- | --- | --- | --- | --- | --- | --- | --- | --- | --- | --- |
| Treatment # | 5 | 1 | 6 | 14 | 2 | 3 | 4 | 7 | 8 | 9 | 10 | 11 | 12 | 13 |
| Karim (%) | 100 |  |  |  | 25 | 50 | 75 | 25 | 50 | 75 |  | 25 | 50 | 75 |
| Salim (%) |  | 100 |  |  | 75 | 50 | 25 |  |  |  | 50 | 37.5 | 25 | 12.5 |
| Monastir (%) |  |  | 100 |  |  |  |  | 75 | 50 | 25 | 50 | 37.5 | 25 | 12.5 |
| INRAT100 (%) |  |  |  | 100 |  |  |  |  |  |  |  |  |  |  |

| Year 2 |  | Pure stands |  |  | Two-way mixtures |  |  |  |  |  |
| --- | --- | --- | --- | --- | --- | --- | --- | --- | --- | --- |
| Treatment # | 1 | 2 | 3 | 7 | 4 | 8 | 6 | 5 | 9 | 10 |
| Karim (%) | 100 |  |  | 50 | 75 | 87.5 | 50 | 75 | 87.5 |  |
| INRAT100 (%) |  | 100 |  | 50 | 25 | 12.5 |  |  |  | 50 |
| Monastir (%) |  |  | 100 |  |  |  | 50 | 25 | 12.5 | 50 |

For the second year the mixtures were changed, based on the first year’s data. Salim was not included, but INRAT100 was used in mixtures with Karim. Karim was mixed with the two cultivars in one-way mixtures in proportions 0%, 50%, 75%, 87.5% and 100% (excluding 25% but adding 87.5%, compared to the first year). There were 10 treatments in total: three pure stands, six one-way mixtures with Karim, and one 50-50 mixture of the resistant cultivars (Table 1). Each treatment was replicated four times in different plots. In the second year, all treatments were also replicated four times without inoculation but with fungicide sprays. This allowed us to have disease-free controls (see next section).

### Preparation of plant and fungal material

Seeds of different cultivars were mixed in the drill according to the proportions of seed numbers corresponding to each treatment. Seed numbers were adjusted by weighing the seeds and using thousand-kernel-weight (TKW) of each cultivar. The seeds were sown on December 27, 2017 (first year), on plots of 5.25 m^2^ (3.5 m x 1.5 m) with a precision seeder at a density of 400 seeds m^-2^, and on November 8, 2018 (second year) on plots of 6 m^2^ (5 m x 1.2 m) at 350 seeds m^-2^ (densities comparable to common practices in the region). Subsequently, mean density of spikes was found to be 381 spikes m^-2^ in the first year and 297 spikes m^-2^ in the second year in inoculated plots and 300 spikes m^-2^ in the second year in fungicide-treated plots. There were no differences in spike densities between treatments in the first (Kruskal-Wallis p=0.93) or the second year (p=0.40), nor between inoculated and fungicide-treated plots (p=0.73). Spatial arrangement of treatments and replicates is shown in Figure 1a.

Standard agronomic practices were used to ensure adequate crop development. During the first year, herbicide treatment with a mixture of Traxos (1.2 L ha^-1^, Syngenta) and Zoom (180g ha^-1^, Syngenta) was applied on January 25, 2018, at tillering stage corresponding to GS13 [according to Zadoks scale (Zadoks et al., 1974)]. Manual and mechanical weeding was performed on March 19, 2018, (corresponding to stem elongation). Nitrogen fertilizer ammonium nitrate was applied three times at rates of 120 kg ha^-1^, 150 kg ha^-1^ and 150 kg ha^-1^, on February 1, February 25, and April 1, 2018, respectively. No fungicides or insecticides were applied.

During the second year, before sowing, the Roundup herbicide (Monsanto; active ingredient: Glyphosate 360g L^-1^) was applied with a rate of 3L ha^-1^. Manual weeding was done on March 25, 2019. Nitrogen fertilizer ammonium nitrate was applied three times at a rate of 150 kg ha^-1^: on December 20, 2018 (GS30), February 14, 2019 (GS32) and March 25, 2019 (GS40). No insecticides were applied. Five fungicide treatments were applied for the treated-control plots to ensure little or no disease. The first fungicide application was done on January 22, 2019 (Product Opus, Stima; Epoxiconazol 125g L^-1^, 0.75L ha^-1^), the second on February 27, 2019 (Cherokee, Syngenta; Cyproconazole 40g L^-1^, Propiconazol 62.5g L^-1^ and Chlorothalonil 375g L^-1^, 1.5L ha^-1^), the third on March 14, 2019 (Opus, Stima; 0.75L ha^-1^), the fourth on April 11 (Ogam, Stima; Epoxiconazol 125g L^-1^ and Kresoxim-methyl 125g L^-1^, 0.7L ha^-1^) and the fifth on May 3, 2019 (Ogam, Stima; 0.7L ha^-1^).

The inoculum of *Z. tritici* was produced in the wheat Septoria platform laboratory according to Ferjaoui et al. (2015) with slight modifications. Six isolates were obtained from infected leaves of durum wheat collected in the same region and used to prepare the inoculum. The isolates were grown for 6 to 8 days on potato dextrose agar. Inoculum was prepared in 250 mL of yeast-glucose liquid medium (30 g of glucose, 10 g of yeast in 1 L of water). The flasks were inoculated with fresh pieces of *Z. tritici* colonies from agar plates and incubated in a rotary shaker at 100 rpm, at 15°C for 5-7 days. The inoculum concentration was adjusted to 10^6^ spores mL^-1^ and the resulting spore suspension was supplemented with 0.1 % of Tween 20 (Merck, UK) prior to inoculation in the field. Approximately 700 mL of the spore suspension was applied per plot using a sprayer (Efco AT800, Italy). In the first year, wheat plants were inoculated three times, on February 27, March 9, and March 20 2018, corresponding to tillering stage (from GS13 to GS26). During the second year, similarly three inoculation were performed on December 12, December 26, and January 10 (GS13 to GS26).

### Disease assessment

Disease levels were assessed two times in both years: at *t*_1_ on flag-1 leaves (the leaf below the flag leaf) and at *t*_2_ on flag leaves [first year *t*_1_ on 22 April (GS 61) and *t*_2_ on 9 May (GS 75); second year *t*_1_ on April 17 (GS 73) and *t*_2_ on April 25 (GS 75)]. From each inoculated plot, 24 leaves were collected without considering their infection status and in a sparse uniform random manner. The collector bias was minimized by a stringent collection protocol: first, a spike was chosen at random without looking at leaves and avoiding edges of the plot. Only after that selection, the leaf below this spike was collected. During the collection, the leaves were placed in paper envelopes and kept on ice in a thermo-insulated box. After collection, the leaves were taken to the lab and kept at 4-5°C for one to three days before inspection. The leaves were then inspected visually for the presence of pycnidia as a sign of STB infection. The absence of pycnidia was interpreted as absence of STB, even if necrotic lesions were visible. In this way, STB incidence was estimated as the proportion of infected leaves in each plot. After visual examination, the infected leaves were mounted on paper sheets and scanned, as described by Karisto et al. (2018). The uninoculated fungicide-treated plots were inspected visually and confirmed to be healthy.

Scanned leaf images were analysed using ImageJ software (Schindelin et al., 2015) with the help of the automated image analysis macro originally developed by Stewart and McDonald (2014) and further improved by Stewart et al. (2016) and Karisto et al. (2018). The program quantified necrotic leaf area and counted pycnidia on each infected leaf as measures of conditional severity of STB, i.e., the severity only on infected leaves. From this raw data, three quantities were computed: (i) the percentage of leaf area covered by lesions (PLACL); (ii) the density of pycnidia per cm^2^ of lesion area (ρ_lesion_); and (iii) the density of pycnidia per cm^2^ of total leaf area (ρ_leaf_). The three quantities characterise different aspects of conditional severity of STB. PLACL measures the damage induced by the pathogen to the host, ρ_lesion_ measures the degree of pathogen reproduction within the host, and ρ_leaf_ is the product of PLACL and ρ_lesion_ that incorporates both host damage and pathogen reproduction (Karisto et al., 2018). To estimate the full, unconditional severity, each of the three measures of conditional severity was multiplied by disease incidence of the corresponding plot.

At each of the four time points, 200 leaf images were selected to test the accuracy of the automated image analysis. They were first inspected qualitatively for lesion and pycnidia detection errors (i.e., substantial underestimation or overestimation). After excluding erroneous leaves, the accuracy of automated counting of pycnidia was evaluated by counting pycnidia manually on 20 leaves at each time point and comparing the manual counts to the estimates from the image analysis. The leaves for both qualitative and quantitative evaluation were selected with a stratified random sampling based on 10 approximately equally sized classes for pycnidia counts to ensure a satisfactory coverage of the entire distribution of pycnidia counts. To quantify the accuracy of automated pycnidia counts with respect to manual counts, we calculated the concordance correlation coefficient (Lin 1989), Pearson’s and Spearman’s correlation coefficients, and the average error of automated pycnidia counts.

### Yield assessment

In all plots, the grain yield (kg ha^-1^) and the TKW (g) were measured. When the plants in all plots had reached physiological maturity (Zadoks et al., 1974), the entire plot area was harvested (29 June 2018 and 24 June 2019). Grain yield was calculated by determining the total grain weight in the harvested area (kg ha^-1^). TKW was estimated by weighing 500 kernels. The effect of disease on yield was determined in the second year’s data by subtracting the yield of each inoculated plot from the yield of the corresponding fungicide-treated plot.

### Statistical analysis

Treatments were compared to each other based on the three measures of the full STB severity described above: PLACL, ρ_lesion_, and ρ_leaf_. From now on, we will use these three measures of severity to refer to full severity, unless specified otherwise. The data were analysed using the Python programming language (version 3.5.2, https://www.python.org), using the open-source packages SciPy (version 1.0.1; Jones et al., 2001), NumPy (version 1.14.3) and matplotlib (version 2.2.3). The source code developed for the analysis is available from the Dryad Digital Repository (DOI: 10.5061/dryad.r2280gbb5).

Presence of global differences between medians of treatments was tested by the Kruskal-Wallis test implemented as scipy.stats.kruskal function. Post-hoc comparisons were conducted with multiple Dunn’s tests with the false discovery rate correction (Benjamini-Hochberg) using the library scikit_posthocs (version 0.3.8). Separate comparisons were first conducted for each of four time points. Due to the limited number of replicates, strong conclusions cannot be drawn from individual treatments and time points. Hence, after performing comparisons within individual mixtures separately, all data were normalised based on the mean of the susceptible treatment for each variable within the time point, and then pooled based on the proportion of the susceptible cultivar Karim in the mixture. The analysis of the pooled data was conducted in a similar fashion as the analysis within the individual treatment groups described above. The aim of this analysis was to identify general patterns of how disease levels depend on the proportion of a resistant cultivar in the mixture.

Finally, we tested for mixture effects using the following procedure. Expected levels of disease and yield were calculated for each mixture based on the average of the two pure stands (100% susceptible and 0% susceptible), weighted according to their proportions in the mixture (linear expectation). Note that in three-way mixtures the resistant cultivars were always in equal proportions and thus 0% susceptible corresponds to 1:1 mixture of the resistant cultivars Salim and Monastir (Table 1). The deviations from the linear expectation were calculated for each plot. Then the treatments were pooled based on the percentage of the susceptible cultivar as above and Wilcoxon signed rank test (scipy.stats.wilcoxon) was used to determine if those deviations from linear expectation were symmetric around zero. Comparison of treatments with respect to yield was based on TKW and grain yield.

## Results

### Overview of the dataset

Septoria tritici blotch severity was assessed at two time points on all 42 plots in the first year and 40 plots in the second. In total, approximately 3900 leaves were inspected for the presence of the disease. Total number of infected leaves included in the image analysis for measuring severity was 3074 (646 leaves at t_1_ 2018; 638 at t_2_ 2018; 887 at t_1_ 2019; 903 at t_2_ 2019). Based on the qualitative check, 70 leaves were excluded from the dataset. The most common reasons for exclusion were overestimation of lesion area due to insect damage and overestimation of pycnidia counts due to dirt on leaves. Overall, the pycnidia counts by image analysis were consistent with the manual counting in the test set of 80 leaves (106 000 pycnidia detected by macro vs 98 000 manually counted, concordance correlation coefficient = 0.98; r_spearman_ = 0.96, p = 2 x 10^-43^; Fig. S2). The final data set based on 3004 leaf images is available from the Dryad Digital Repository (DOI: 10.5061/dryad.r2280gbb5).

We observed considerable disease levels at each time point. Both the disease and the yield levels were higher in the second year: mean pycnidia density per unit leaf area (ρ_leaf_) on flag leaves (t_2_) was 24 cm^-2^ in 2018 and 103 cm^-2^ in 2019 (p = 6 x 10^-9^), grain yield in 2018 was 2500 kg ha^-1^ vs 5600 kg ha^-1^ in 2019 (p = 7 x 10^-15^, Fig. 2a, b). Disease levels were generally higher in the pure stands of the susceptible cultivar (Karim) than in the pure stands of the resistant cultivars (Salim, Monastir and INRAT100) or mixtures, consistently between all time points (see supplementary figures for different variables and time points: Figs. S3-S4 for pure stands, Figs. S5-S8 for individual mixtures). Overall degree of disease control in mixtures relative to susceptible pure stands in shown in Figure 2d, e.

**Figure 2.**
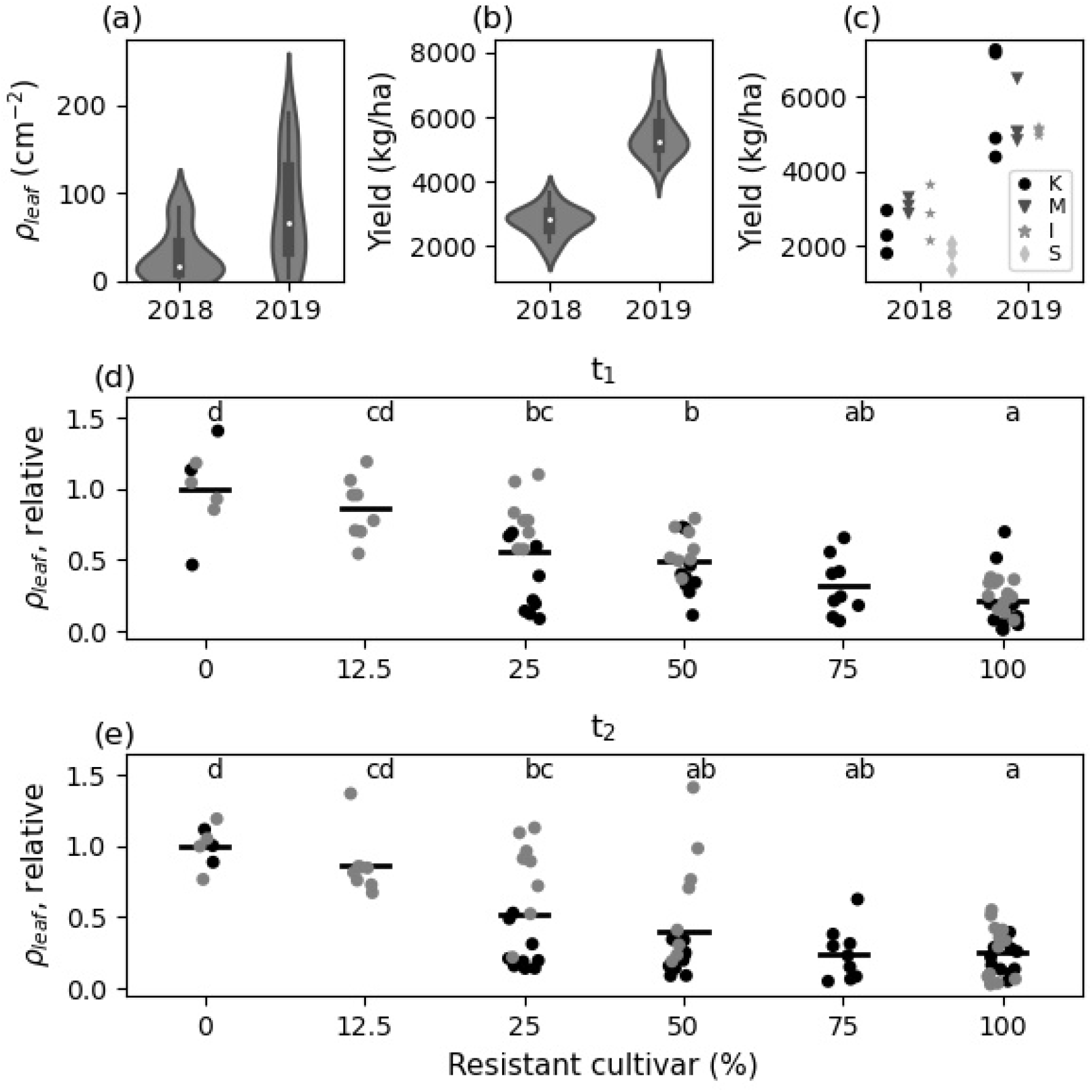
STB severity and yield patterns. Within the grey violin plots in (a) and (b), box plots in black and medians in white. Core of the box plots range from the first to the third quartile, and the whiskers span 1.5 interquartile range further. (a) Distribution of STB severity (ρ_leaf_, plot means) in the repeated treatments in the first year (2018) and the second year (2019). Disease levels were higher in the second year (p=0.015, for repeated treatments). (b) Grain yield in the repeated treatments in both years. Yield was higher in the second year (p=5.7 x 10-7). (c) Grain yield in pure stands. No differences between cultivars within each year (p=0.10 and p=0.87 for the first and the second year, respectively). Letters in legend refer to first letter of the cultivar name. (d, e) Pattern of STB severity at t_1_ and t_2_, respectively (ρ_leaf_, plot means) in the combined data of two years (black: 2018, grey: 2019), including all treatments, pooled based on the percentage of resistant cultivar. Values relative to the susceptible treatments in each time point. Black bars show mean severity. Mixtures with 0% or 12.5%had no significant difference, but 25%, 50% and 75% had lower severity than 0% (only susceptible) and were not different from each other.

### Disease reduction in the mixtures

Analysis of disease levels in durum wheat mixtures revealed a general pattern where the pure stand of the susceptible cultivar Karim had the highest disease levels and treatments including a resistant cultivar had significantly lower disease levels, except for the smallest proportion of 12.5% of resistant cultivar (Figure 2d, e). Introducing only 25% of a resistant component to the susceptible pure stand was enough to suppress the disease levels significantly by 48% compared to the susceptible pure stands (p = 0.0198; see also Figs. S9-S10 for combined performance of mixtures within each time point). In the first year, mixtures with only 25% of a resistant component suppressed the disease down to levels not significantly different from the resistant pure stand (Fig. S9f). These trends were generally true for all three measures of severity, both two-way and three-way mixtures and the four time points with a few exceptions (see supplementary Figures).

For yield measures, there was neither a clear increase nor decrease when the proportion of the susceptible cultivar Karim was decreased from 100% to zero. A possible explanation for this observation is that in pure stands, the susceptible cultivar Karim did not have consistently lower or higher yield compared to the pure stands of the resistant cultivars (Fig 2c, no significant differences).

### Mixture effects

If each component of a mixture performed independently of other components, then the overall disease severity in a mixed stand would be given by the sum of disease severities in corresponding pure stands weighted by the mixing proportions. However, mixture effects may improve the overall performance compared to the sum of individual components. To determine the magnitude of mixture effects, we compared disease severity and yield observed in mixed stands to linear expectations based on measurements in pure stands. Comparing mixture treatments to linear expectations of each treatment group (In 2018 Karim-Salim, Karim-Monastir, and three-way; in 2019 Karim-Monastir and Karim-INRAT100) showed that adding 25% of resistance to susceptible Karim resulted in significant beneficial mixture effect on disease levels (ρ_leaf_), i.e. the severity was lower than expected (p = 0.0099). The mixture effect was strongest in the treatment with 25% of resistant component (mean effect = −30%-points) but significant also for 50% and 75% of resistance (−23%-points, p=0.0129 and −21%-points, p=0.0284, respectively. Fig 3). See Figure S11 for mixture effect tests on other variables, and Figures S12-S13 for the tests on individual time points separately. Mixture effects were generally stronger and more common in the first year than in the second. We observed a beneficial mixture effect on yield in 25% resistant mixtures in the first year (Fig. S12g) but not in the pooled data of two years (Fig. S11c, d).

**Figure 3.**
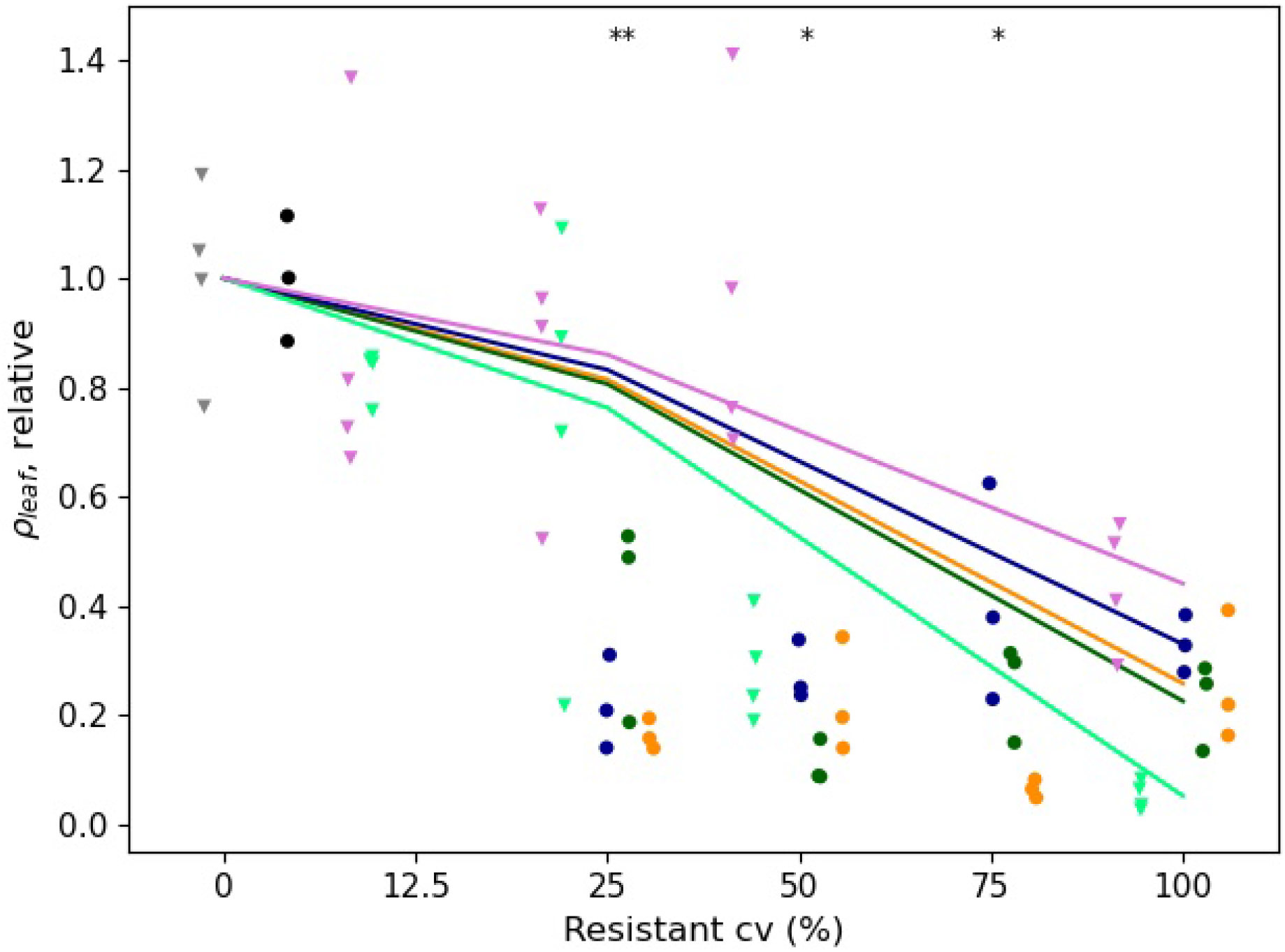
Non-linear mixture effects on STB severity at t2 of both years (ρleaf, plot means, relative to mean of the susceptible treatment). Lines show linear expectations for each group of treatments (note non-uniform scale of the x-axis). Dark blue, yellow and green dots and lines represent Karim-Salim, Karim-Monastir and three-way mixtures, respectively, in the first year. Light green and pink triangles and lines represent Karim-Monastir and Karim-INRAT100 mixtures, respectively, in the second year. Black dots: susceptible cultivar Karim in the first year, grey triangles: Karim in the second year. Significant mixture effects, i.e. lower severity than expected, are indicated with asterisks: * for p<0.05 and ** for p<0.01.

### Effect of the number of resistant components in the mixture

To determine whether adding a second resistant cultivar to a two-way mixture of a resistant and a susceptible cultivar leads to a further reduction of disease, we compared the STB severity and yield in two- and three-way mixtures at a constant proportion of the susceptible cultivar Karim (Figs. S14-S17). For example, at 75% of Karim, we compared the three-way mixture of 75%/12.5%/12.5% proportions of Karim/Salim/Monastir with each of the corresponding two-way mixtures 75%/25% Karim/Salim and 75%/25% Karim/Monastir. In some cases, three-way mixtures had a slightly higher STB severity than both corresponding two-way mixtures; in other cases, three-way mixtures had an intermediate severity with respect to corresponding two-way mixtures. Interestingly, in none of the cases, a mixture with two resistant components exhibited a significant reduction in STB severity or increase in yield with respect to both corresponding mixtures with only one resistant cultivar.

### Cultivar mixtures combined with fungicides

The fungicide-treated plots yielded more than the inoculated, not fungicide-treated plots. Mean effect on yield was 1500 kg ha^-1^ (SD=790) and on thousand kernel weight 7.4 g (SD=5.7). There were no differences between treatments on the effect of fungicides for either measure of yield. Moreover, when pooling the treatments based on percentage of resistant cultivar, we found no significant differences (Fig. S18). All fungicide-treated plots were fully protected from fungal diseases by the intensive spray program applied.

## Discussion

Our analyses show that introducing only 25 % of a disease-resistant cultivar into a pure stand of a susceptible cultivar provides a substantial reduction in disease levels. To measure the severity of Septoria tritici blotch in the field, we used a high-throughput method based on digital analysis of leaf images. This method allowed us to acquire a large dataset that exceeds previous studies on host mixtures with respect to accuracy and precision. In addition, this method is an improvement particularly when studying mixtures that have an increased variance in disease levels and morphology among plants in mixtures compared to single genotype plots, which can make visual assessments more challenging. Finally, the method allowed us to accurately measure numbers of pycnidia on wheat leaves, thereby allowing us to quantify pathogen reproduction within the host, which was not possible using conventional visual assessment.

The observed pattern matches qualitatively with the findings of Jeger et al. (1981b), who investigated the effect of cultivar mixtures on the epidemic development of Septoria nodorum blotch (SNB) caused by the necrotrophic fungal pathogen *Parastagonospora nodorum* (Oliver et al., 2012; Quaedvlieg et al., 2013). Jeger et al. (1981b) mixed an SNB-resistant with an SNB-susceptible cultivar at different proportions and reported that the 25%/75% resistant/susceptible mixture “reduced disease levels effectively to that found in the resistant pure stand” – similarly to what we observed in the first year. This similarity in outcomes for *P. nodorum* and *Z. tritici* suggests that there may be a general underlying mechanism. Since the two pathogens are somewhat similar in terms of their epidemiology, infection biology and population genetic structure, further studies on cultivar mixtures affecting these and other similar pathogens could establish whether this pattern holds more generally and to determine which characteristics of the pathogens are responsible for this effect.

Short-range splash dispersal that dominates the spread of both *P. nodorum* and *Z. tritici* may be one such characteristic. The disease-reduction pattern described above may be favouring the barrier effect rather than the dilution effect as the dominant mechanism of disease reduction. Note however, that the pattern was nearly linear in the second year, favouring dilution. This is because the dilution effect is expected to cause a gradual decrease in the level of disease when the proportion of resistant plants is increased, as predicted for example by the discrete-time population model of Jeger et al. (1981a). In contrast, the barrier effect may result in an abrupt, threshold-like drop in the level of disease at a certain critical proportion of the resistant cultivar in the mixture. At this critical proportion, the connectivity between susceptible plants is disrupted and their population is subdivided into isolated patches. Our data from the second year suggests that 12.5% of the resistant component in the mixture is not enough for achieving this critical proportion and substantial reduction of the disease.

Similar fragmentation phenomena have been studied extensively in ecology in the context of habitat loss and fragmentation by adapting the conceptual framework of percolation theory (Bascompte and Sole, 1996; Swift and Hannon, 2010), but have not received attention in studies of cultivar mixtures. This mechanism is expected to be of importance in pathogens with sufficiently short-range dispersal (for example, splash dispersal in *Z. tritici* and *P. nodorum* on wheat and *Rhynchosporium secalis* on barley, or dispersal of many soil-borne pathogens), but to be less prominent with air-borne pathogens such as those causing rust and mildew diseases. Theoretical prediction of the critical proportion of resistant plants that causes fragmentation of the susceptible plant population requires quantitative characterisation of a pathogen’s dispersal under field conditions. Such measurements in *Z. tritici* and *P. nodorum* are still largely lacking, although recently, Karisto et al. (2019b) estimated dispersal kernels of *Z. tritici* in the field.

Adding a second resistant cultivar to a two-way mixture making it a three-way mixture provided no additional reduction of disease compared to two-way mixtures of resistant and susceptible cultivars, contrary to the theoretical prediction (Mikaberidze et al., 2015). A possible explanation for this discrepancy could be that (i) the pathogen population did not possess a sufficient degree of specialisation with respect to the two resistant cultivars and therefore the pathogen population has not been partitioned enough between the two resistant cultivars to result in a measurable reduction of disease. Alternatively, (ii) the resistance of the two cultivars has a largely overlapping genetic basis. Both of these scenarios violate the assumptions of the model (Mikaberidze et al., 2015).

Although we did not observe a significant reduction of disease when a second resistant component was added to two-way mixtures, this can have an important effect on the adaptation of pathogen populations to host resistances. If the resistance in the two resistant cultivars is conferred by different genes, then a three-way mixture is expected to impose less selection on the pathogen population compared to a two-way mixture with the same proportion of the susceptible component, in this way extending the durability of host resistances.

The differences in the performance of cultivar mixtures in disease reduction between years are likely to reflect remarkably differing weather conditions. The second year, 2019, was exceptionally good for wheat yield in Tunisia, which was reflected in our data. These conditions were favourable also for STB leading to higher levels of disease severity in the second year. Possible other sources of variability include length of the growing season and timings of inoculation and leaf collections (Fig. S1). A curious detail in our experiment is that the mixture effects for disease control and yield were stronger in the first year, i.e. the year of lower yield. Similarly, Gigot et al. (2013) observed the susceptible cultivar was consistently protected in mixture under low to moderate disease pressure. If further experiments would confirm consistency of this pattern, the mixtures would be particularly useful for farmers during “bad” years and hence provide a convenient protection against extreme losses. Long-term experiments would be desirable for establishing benefits of cultivar mixtures in variable conditions.

To conclude, our study (together with that of Jeger et al. 1981b) contributes to establishment of a practically useful rule of thumb, according to which adding 25% of resistant plants to the susceptible pure stand provides substantial protection from disease, in the best case as strong as the resistant pure stand. Such mixtures may have an important advantage with respect to planting resistant pure stands: they are more likely to be used by growers if the susceptible cultivar is agronomically superior compared to the resistant cultivar and/or is generally more accepted by growers. A followup study will need to consider whether these mixtures would behave differently or similarly under natural STB infection and in different environments as well as to examine the durability of the disease control provided by a mixture over time.

## Acknowledgements

This work was supported by the “CRP WHEAT Tunisia Septoria Precision Phenotyping Platform”. PK and AM gratefully acknowledge financial support from the Swiss National Science Foundation through the Ambizione grant PZ00P3_161453. The authors have no conflicts of interest to declare.

## Data availability

The data and the source code for statistical analysis are available in Dryad Digital Repository (DOI: 10.5061/dryad.r2280gbb5).

## Notes

https://doi.org/10.5061/dryad.r2280gbb5

### Competing Interest Statement

The authors have declared no competing interest.

### Summary of Updates

We revised the main text, adding some references and removing unnecessary ones. We improved readability of all figures. We added references to the dataset in Dryad digital repository.

