## Supplementary Figures for "Improved control of Septoria tritici blotch in durum wheat using cultivar mixtures"

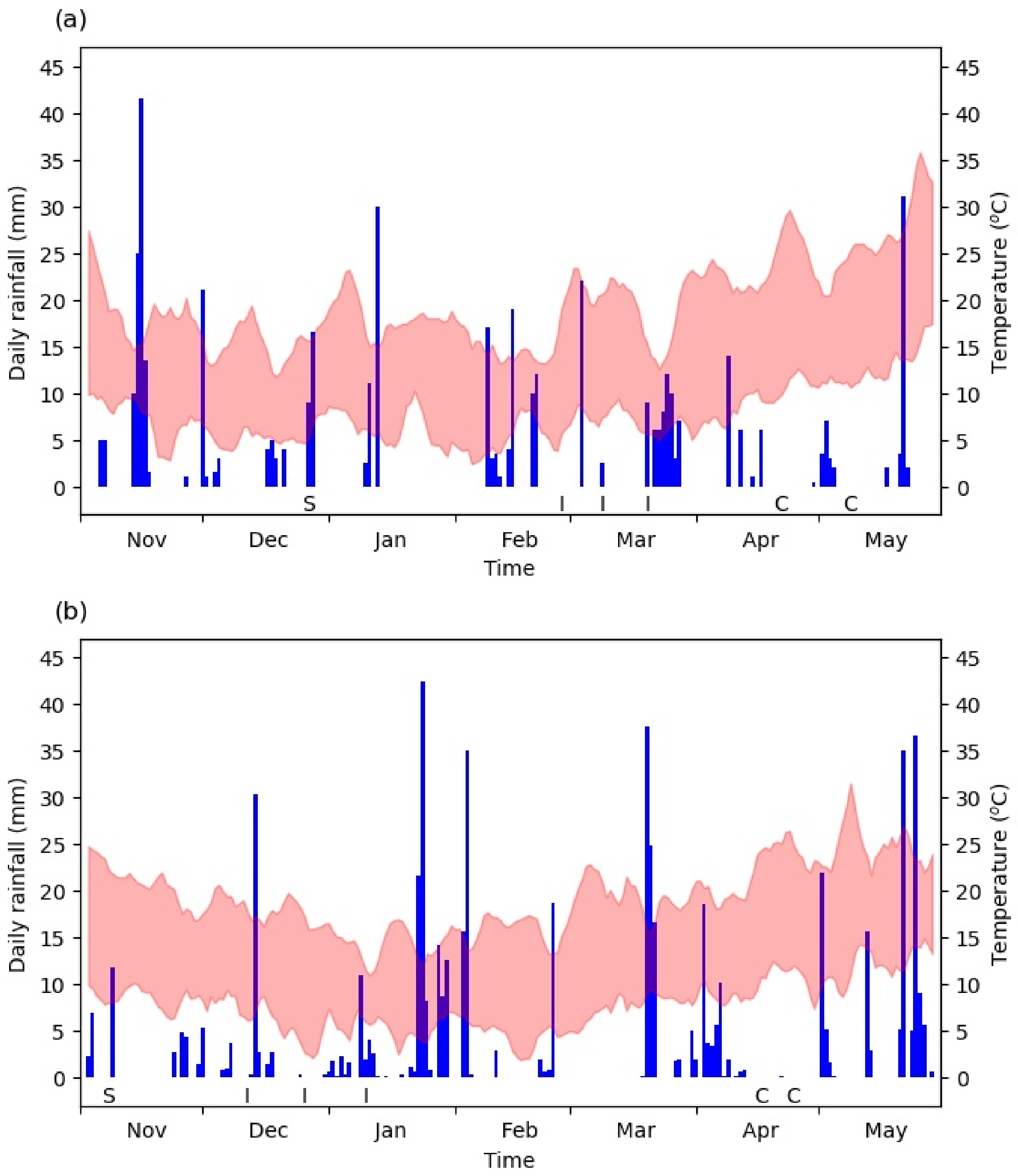


**Figure S1**. Weather conditions during the experiment. Daily precipitation in blue bars and 5-day moving averages of minimum and maximum temperatures in red. Dates of important experimental procedures are indicates by letters: S -- sowing, I -- inoculation, C -- leaf collections for disease assessment. (a) The first year, (b) the second year.

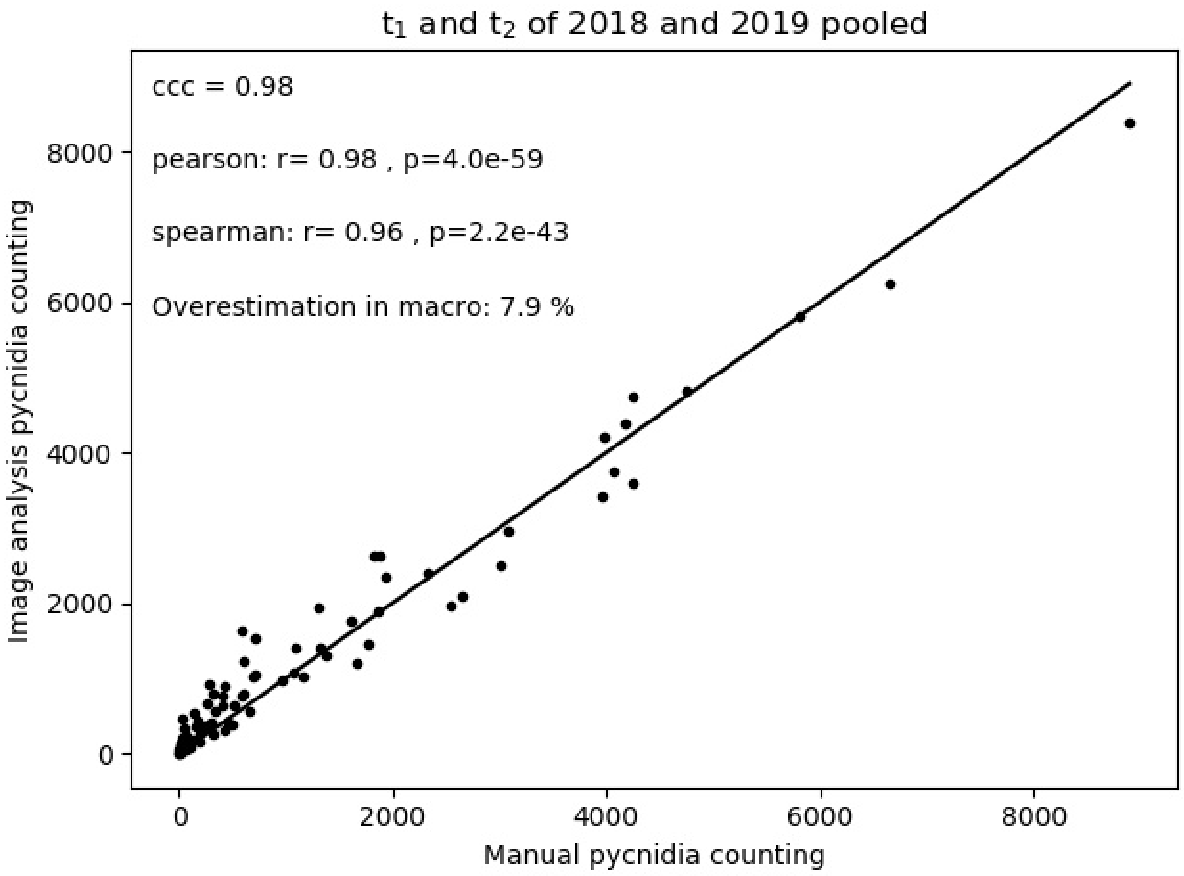


**Figure S2**. Comparison of pycnidia counts between the automated image analysis macro and visual counting on scanned leaves. The solid line represents perfect agreement. Concordance correlation coefficient (ccc) representing the overall agreement of the two measures is 0.98. The mean of the manual counts was about 8% lower than macro counts.

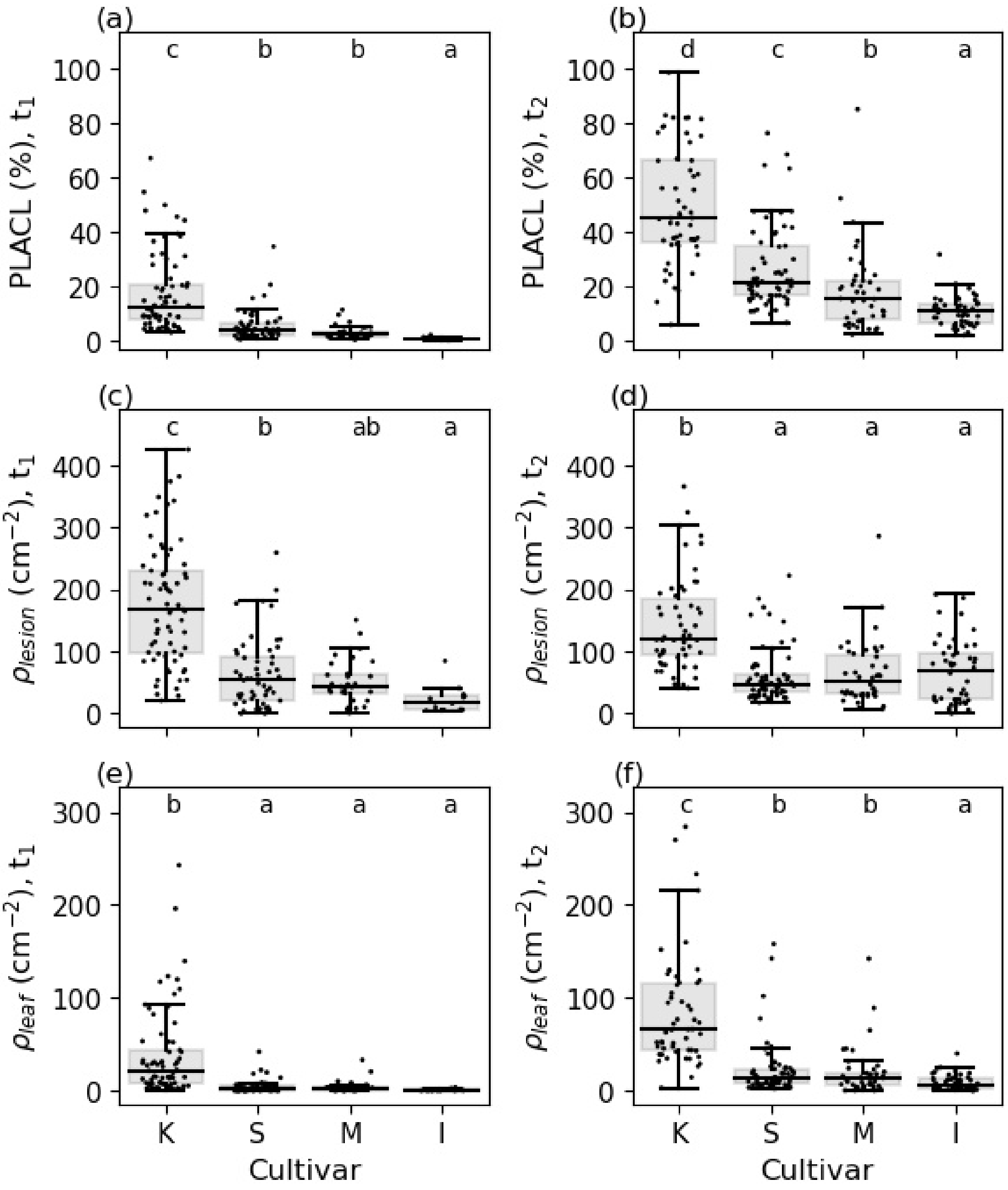


**Figure S3.** Comparison of STB severity in pure stands of the four cultivars in the first year: Karim (K), Salim (S), Monastir (M) and INRAT100 (I). Karim has substantially higher STB severity than the three newer cultivars. (a, b) PLACL, (c, d) $\rho_{lesion}$ (e, f) $\rho_{leaf}$. Left column: $t_1$, right column: $t_2$.

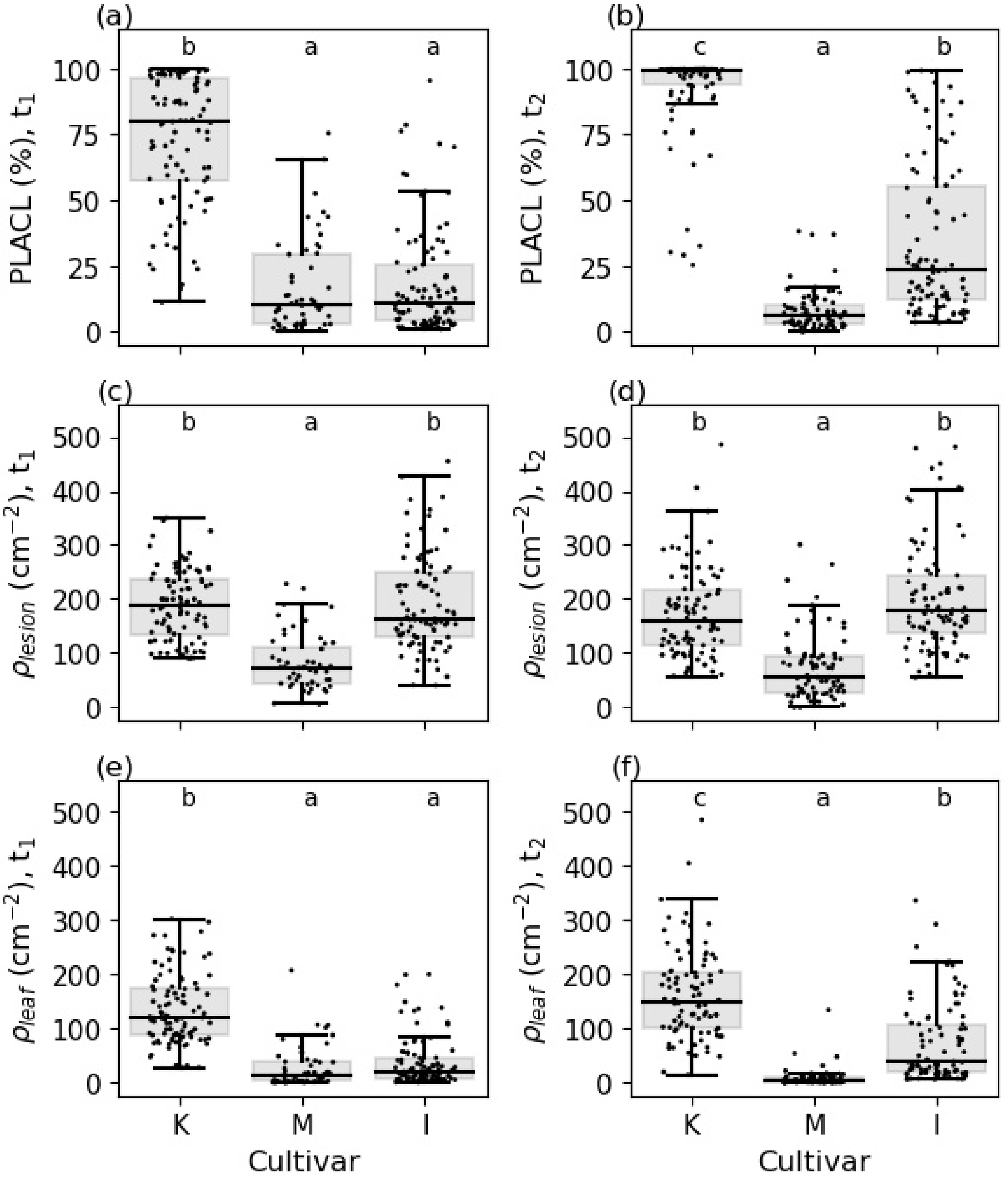


**Figure S4.** Comparison of STB severity in pure stands of the three cultivars in the second year: Karim (K), Monastir (M) and INRAT100 (I). (a, b) PLACL, (c, d) $\rho_{lesion}$ (e, f) $\rho_{leaf}$. Left column: $t_1$, right column: $t_2$.

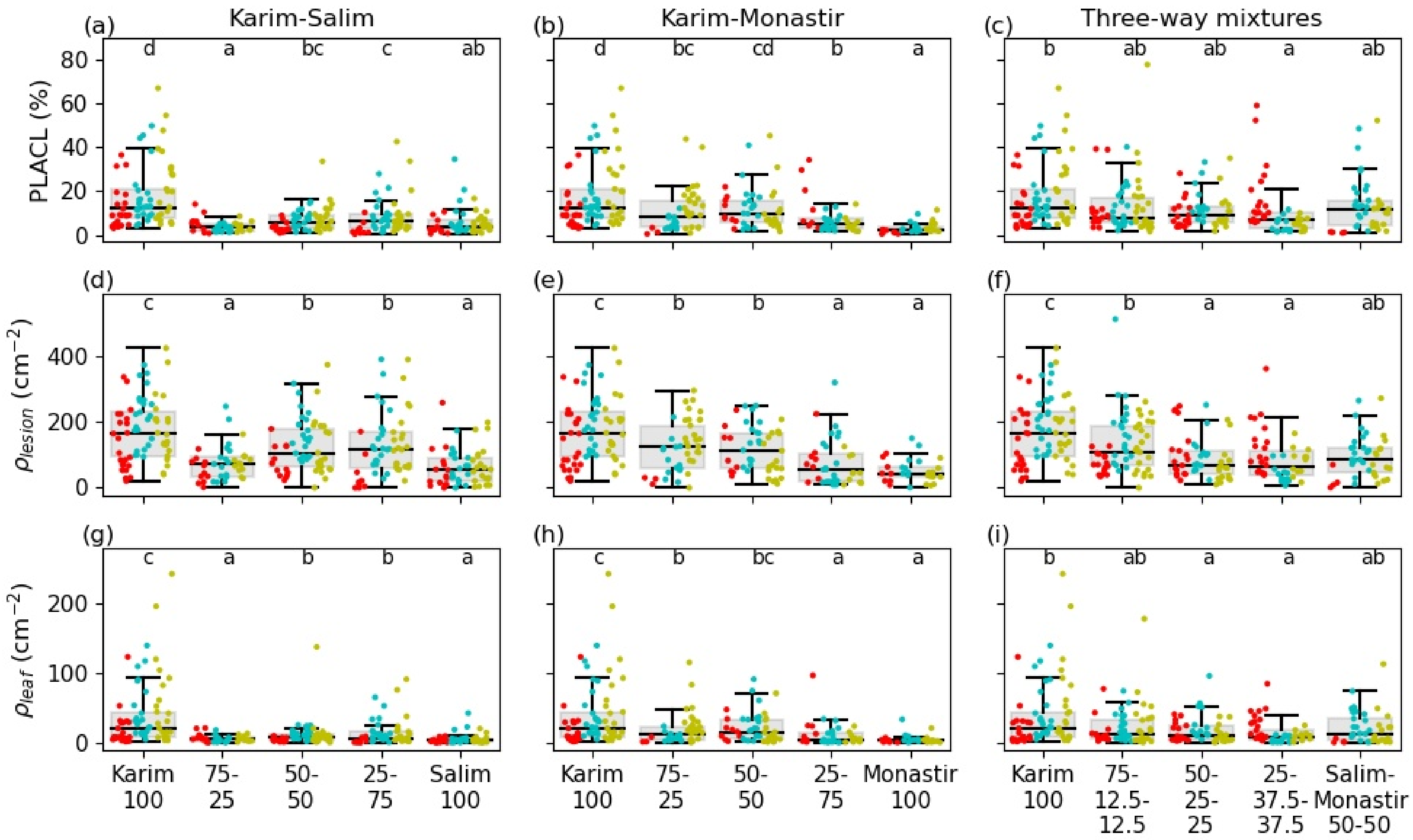


**Figure S5.** STB severity at $t_1$ measured in different mixtures in the first year as PLACL (a-c), $\rho_{lesion}$ (d-f) and $\rho_{leaf}$ (g-j). Letters indicate significant pairwise differences. Red, cyan and yellow: 1st, 2nd and 3rd replicate, respectively.

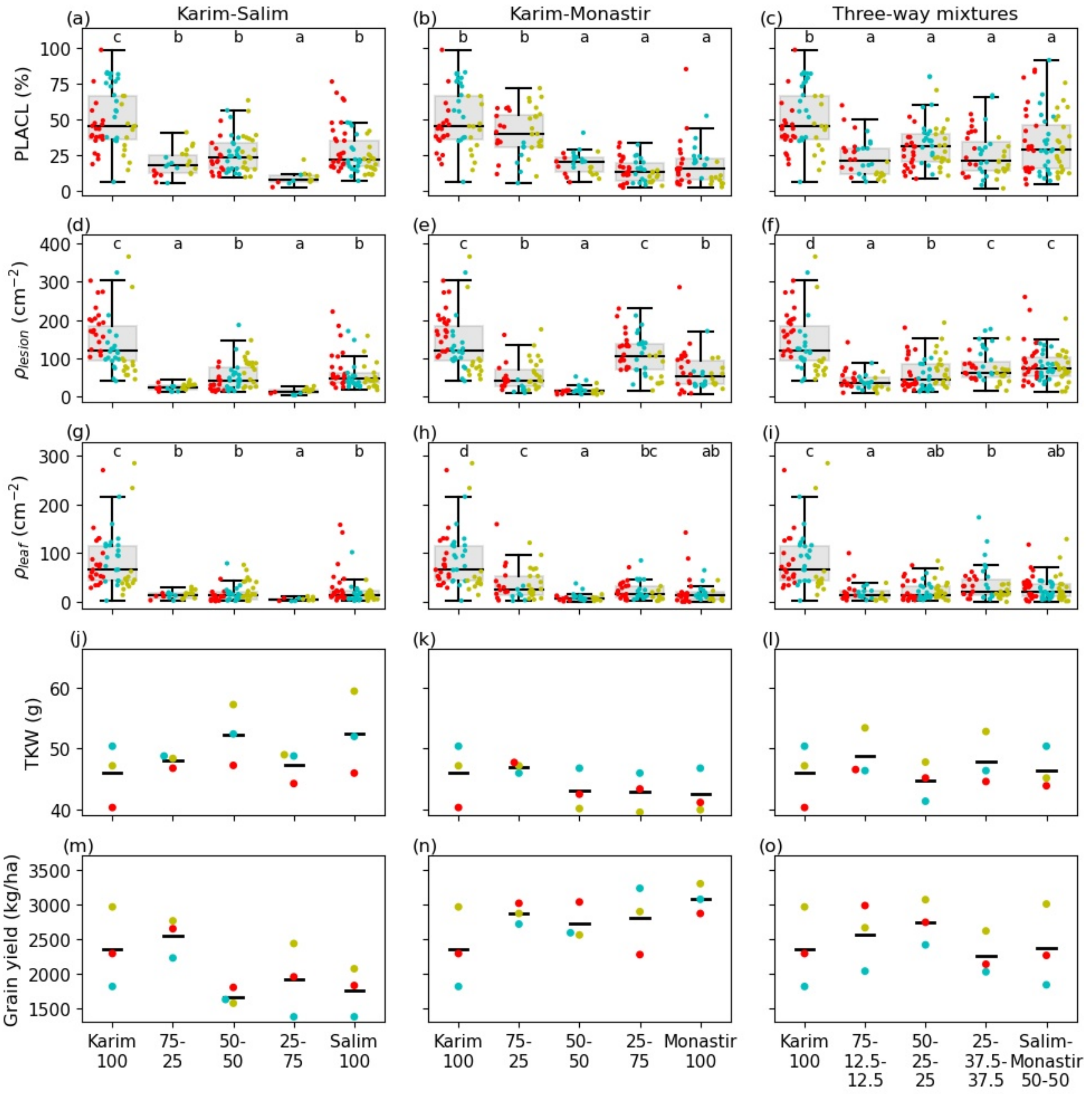


**Figure S6.** STB severity at $t_2$ and wheat yields in different mixtures in the first year. STB severity measured as PLACL (a-c), $\rho_{lesion}$ (d-f) and $\rho_{leaf}$ (g-j); yield measured in TKW (j-l) and in grain yield (m-o). Letters indicate significant pairwise differences. Red, cyan and yellow: $1^{st}$, $2^{nd}$ and $3^{rd}$ replicate, respectively.

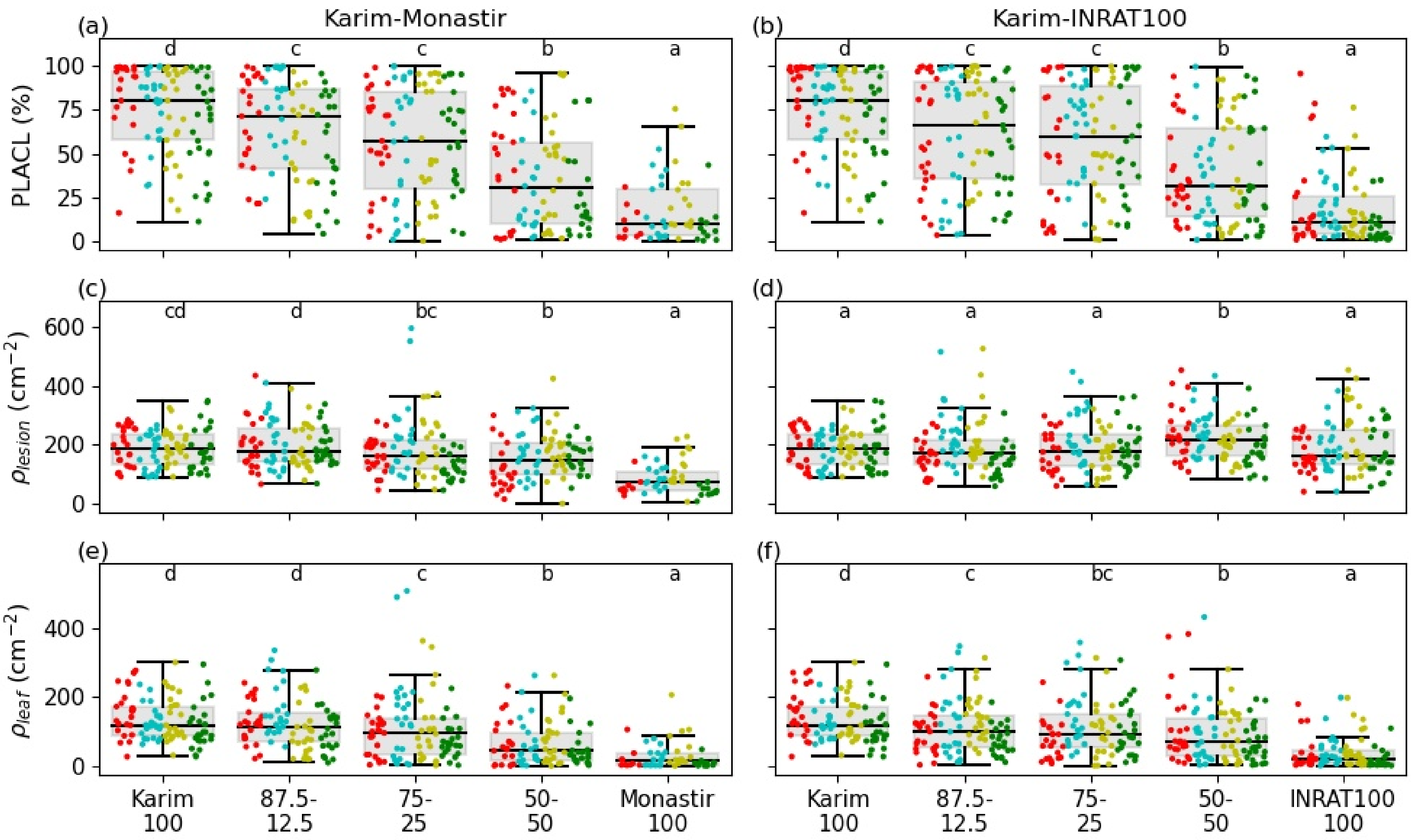


**Figure S7.** STB severity at $t_1$ measured in different mixtures in the second year as PLACL (a-c), $\rho_{lesion}$ (d-f) and $\rho_{leaf}$ (g-j). Letters indicate significant pairwise differences. Red, cyan, yellow and green: 1st, 2nd, 3rd and 4th replicate, respectively.

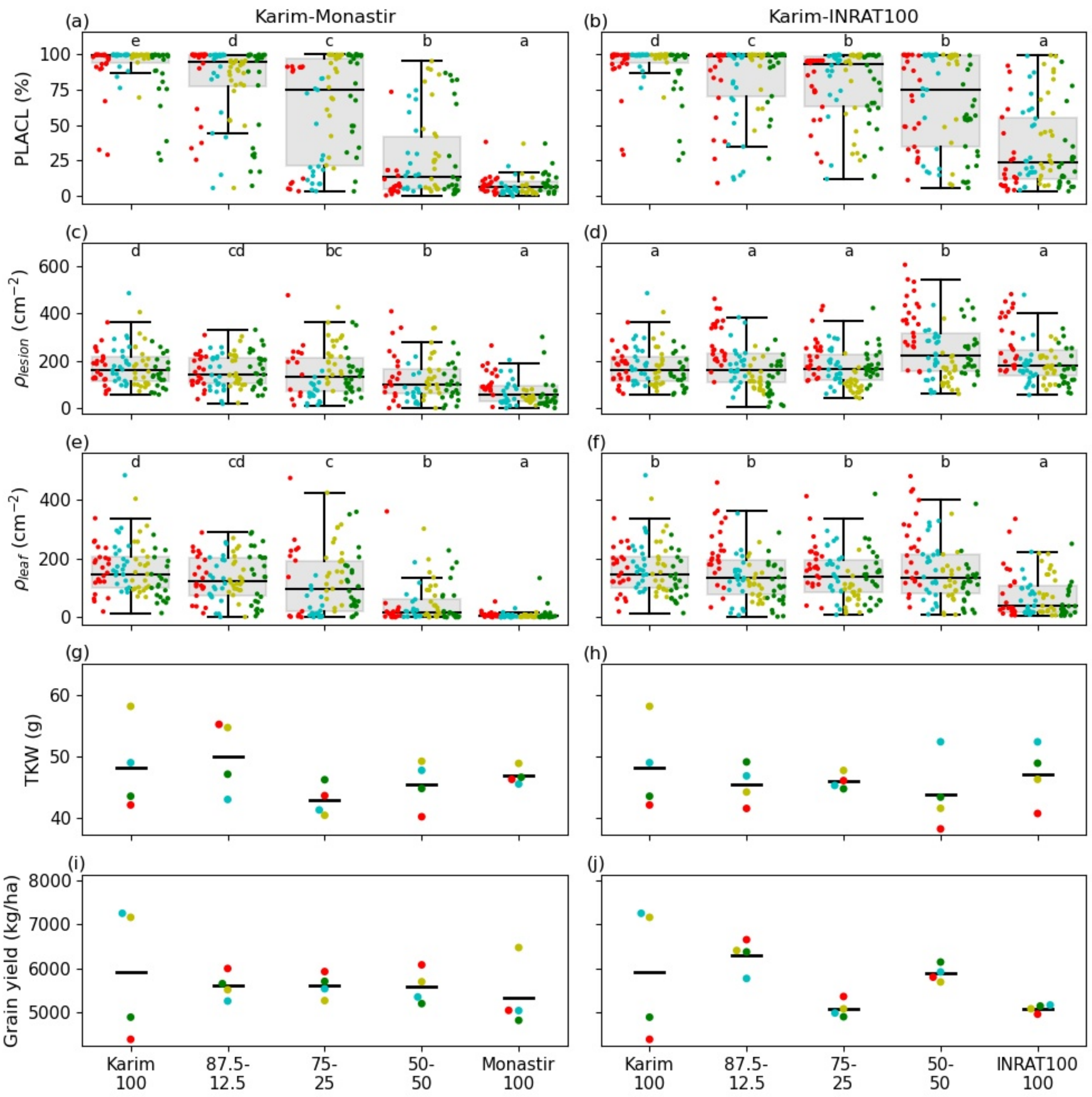


**Figure S8.** STB severity at $t_2$ and wheat yields in different mixtures in the second year. STB severity measured as PLACL (a-c), $\rho_{lesion}$ (d-f) and $\rho_{leaf}$ (g-j); yield measured in TKW (j-l) and in grain yield (m-o). Letters indicate significant pairwise differences. Red, cyan, yellow and green: 1st, 2nd, 3rd and 4th replicate, respectively.

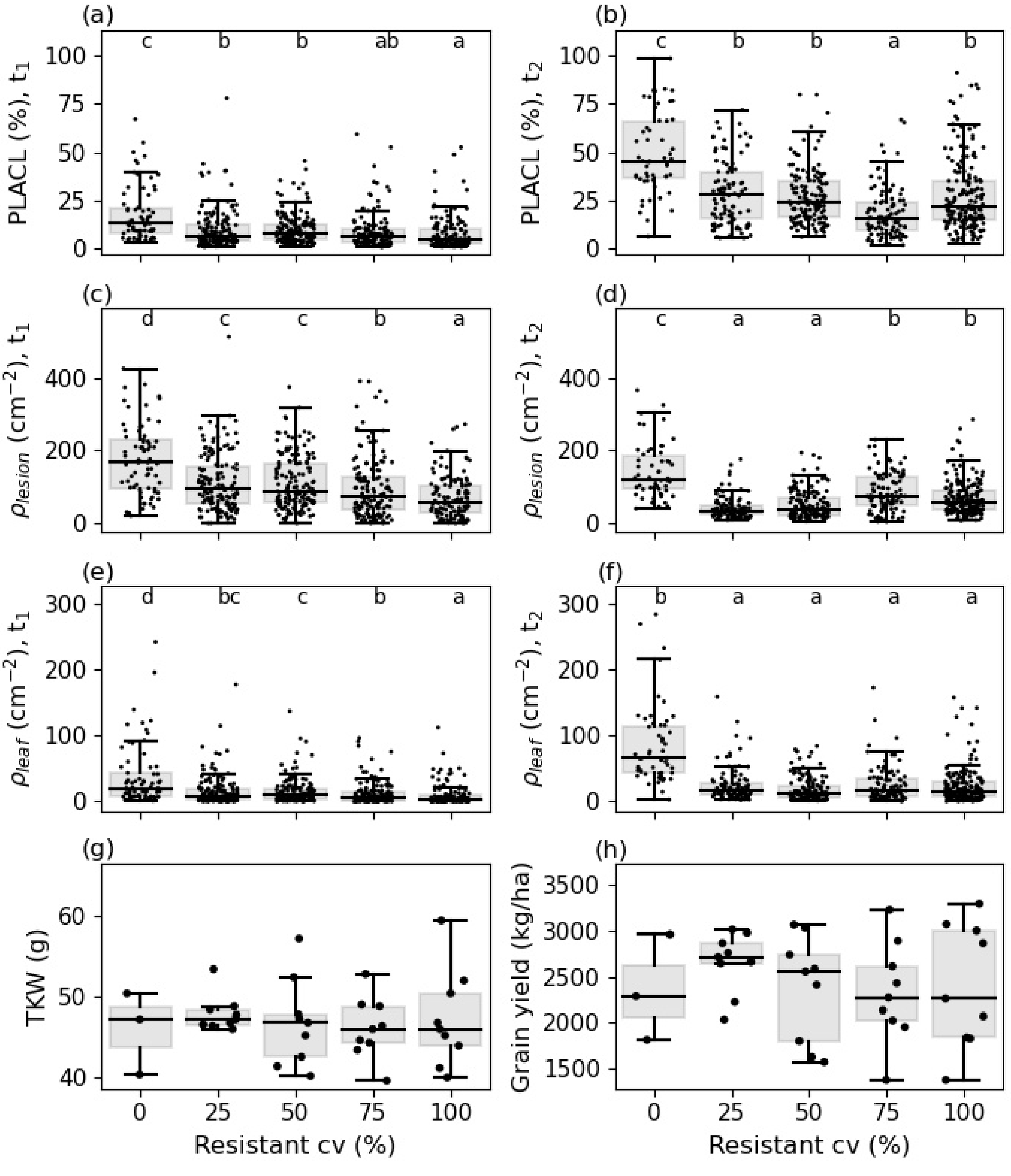


**Figure S9.** Measures of STB severity at $t_1$ (a, c, e) and $t_2$ (b, d, f), and yield (g-h) in the first year. Data from different treatments is pooled based on the proportion of the resistant component in the mixture. Disease severity patterns measured as PLACL (a-b), $\rho_{lesion}$ (c-d) and $\rho_{leaf}$ (e-f) and yield measured in TKW (g) and grain yield (h). No significant differences were detected in yield data.

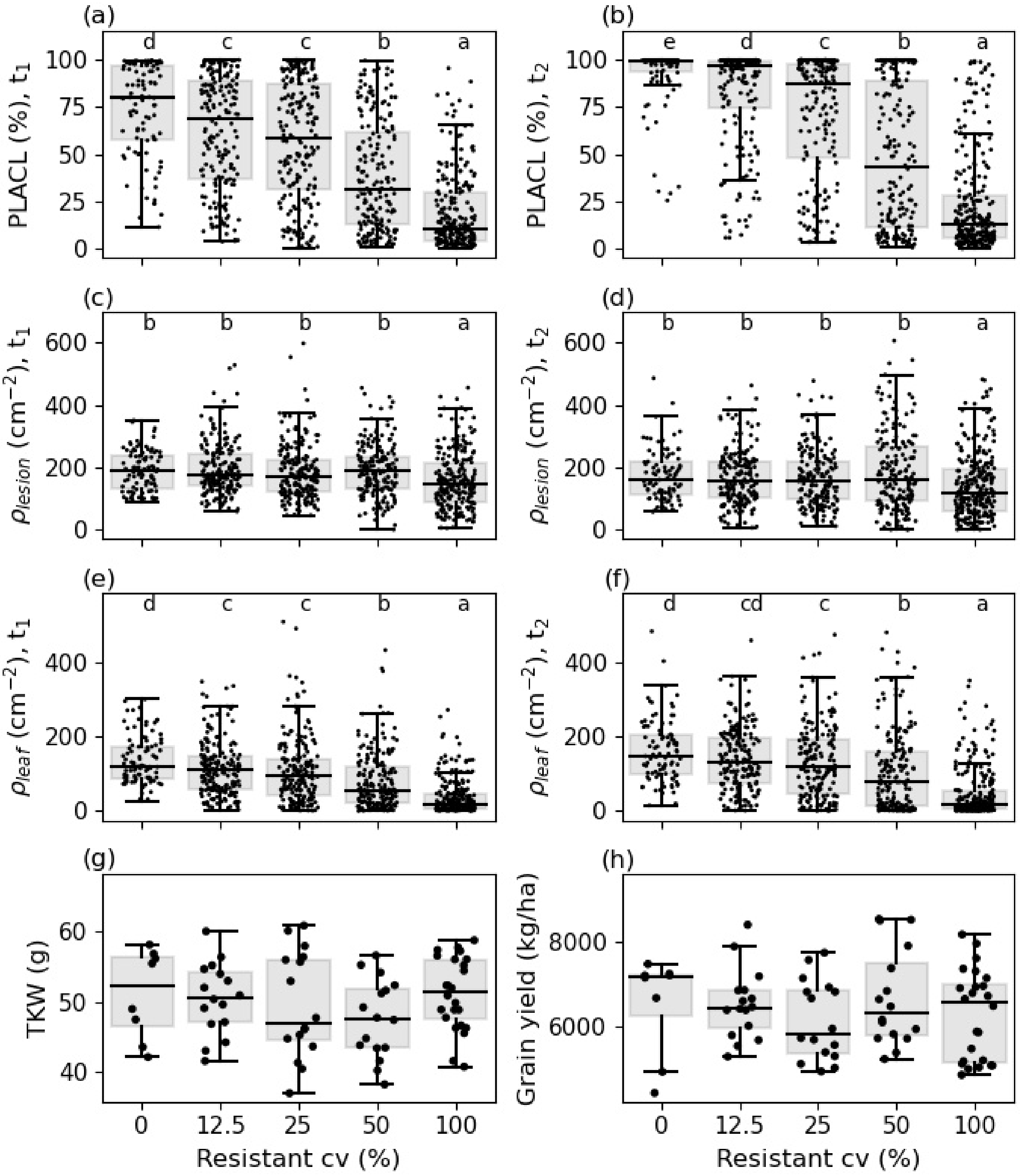


**Figure S10.** Measures of STB severity at $t_1$ (a, c, e) and $t_2$ (b, d, f), and yield (g-h) in the second year. Data from different treatments is pooled based on the proportion of the resistant component in the mixture. Disease severity patterns measured as PLACL (a-b), $\rho_{lesion}$ (c-d) and $\rho_{leaf}$ (e-f) and yield measured in TKW (g) and grain yield (h). No significant differences were detected in yield data.

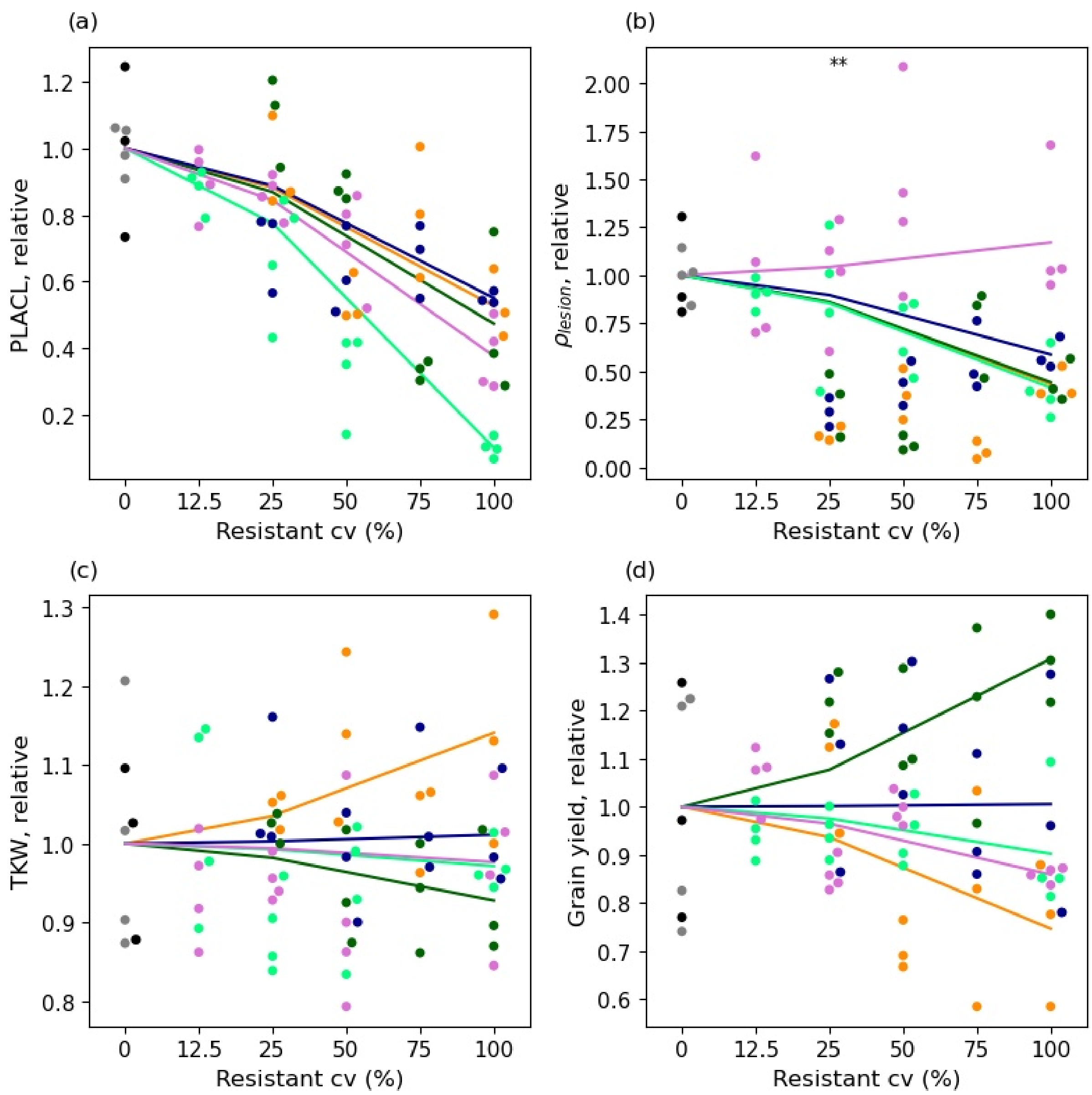


**Figure S11.** Mixture effects on STB severity and yield relative to the mean of susceptible treatment. Structure of the panels is similar to Figure 3, but here we used variables PLACL (a), $\rho_{lesion}$ (b), TKW (c) and grain yield (d). Each point represents one plot and lines show linear expectations for each group of treatments (note non-uniform scale of x-axis). Dark blue, yellow and green dots and lines represent Karim-Salim, Karim-Monastir and three-way mixtures, respectively, in the first year. Light green and pink represent Karim-Monastir and Karim-INRAT100 mixtures, respectively, in the second year. Black and grey dots represent data for the susceptible cultivar Karim in the fist and the second year, respectively. Significant deviations from the linear expectation are indicated with asterisks: * for p<0.05 and ** for p<0.01.

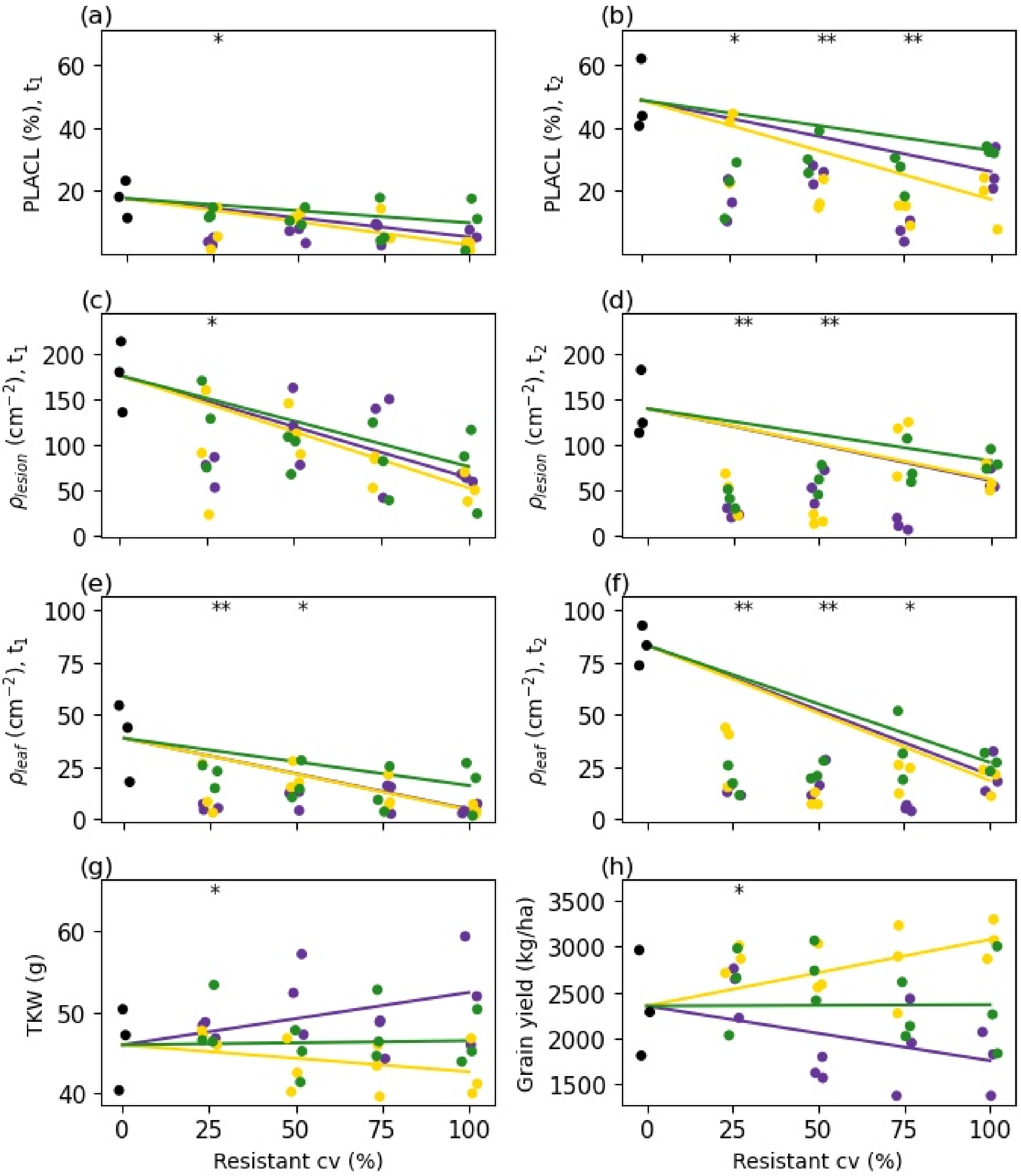


**Figure S12.** Mixture effects on STB severity and yield in the first year. Structure of the panels is similar to Figure 3, but here all variables and both time points are shown: PLACL (a, b), $\rho_{lesion}$ (c, d), $\rho_{leaf}$ (e-f), TKW (g) and grain yield (h). Each point represents one plot and lines show linear expectations for each group of treatments. Purple, yellow and green dots and lines represent Karim-Salim, Karim-Monastir and three-way mixtures, respectively. Black dots represent the susceptible cultivar Karim. Significant deviations from the linear expectation are indicated with asterisks: * for $p<0.05$ and ** for $p<0.01$.

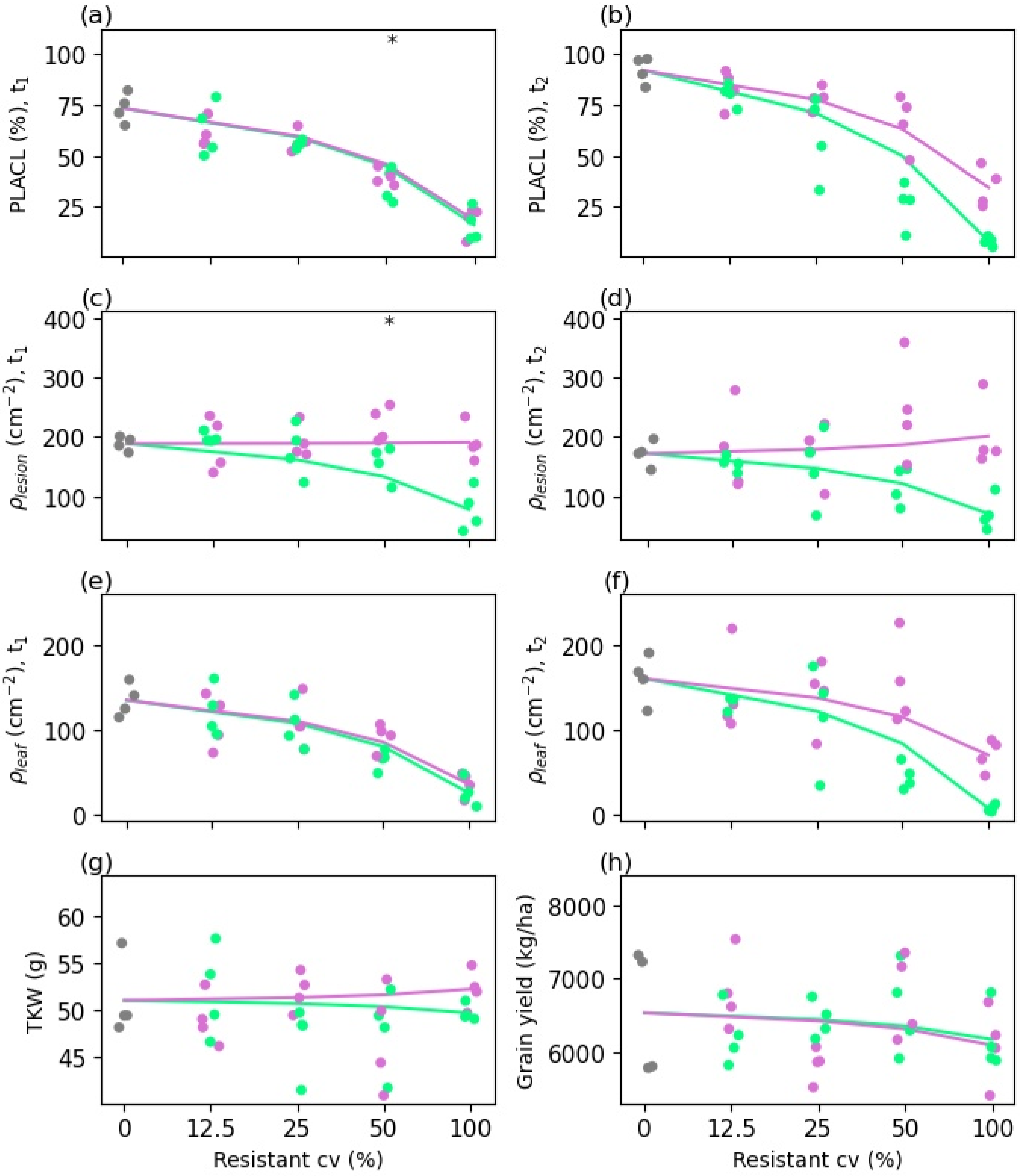


**Figure S13.** Mixture effects on STB severity and yield in the second year. Structure of the panels is similar to Figure 3, but here all variables and both time points are shown: PLACL (a, b), $\rho_{lesion}$ (c, d), $\rho_{leaf}$ (e-f), TKW (g) and grain yield (h). Each point represents one plot and lines show linear expectations for each group of treatments (note non-uniform scale of x-axis). Light green and pink represent Karim-Monastir and Karim-INRAT100 mixtures, respectively, in the second year. Grey dots represent the susceptible cultivar Karim. Significant deviations from the linear expectation are indicated with asterisks: * for p<0.05 and ** for p<0.01.

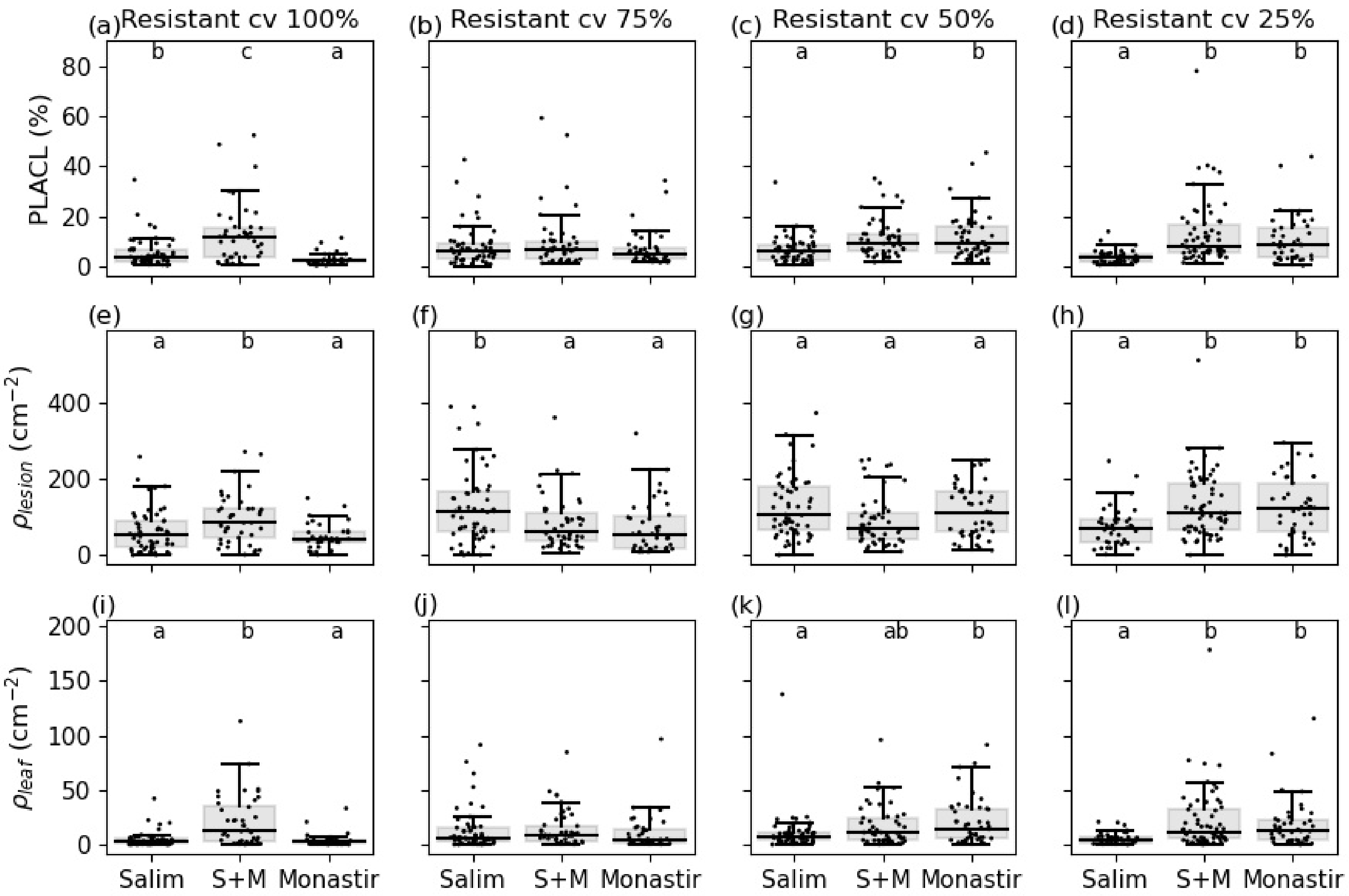


**Figure S14.** The effect of the number of resistant components in the mixture. STB severity at $t_1$ (flag-1 leaves) of the first year. Combinations of the resistant cultivars in equal proportions do not reduce STB severity compared to mixtures with only one of the resistant cultivars.

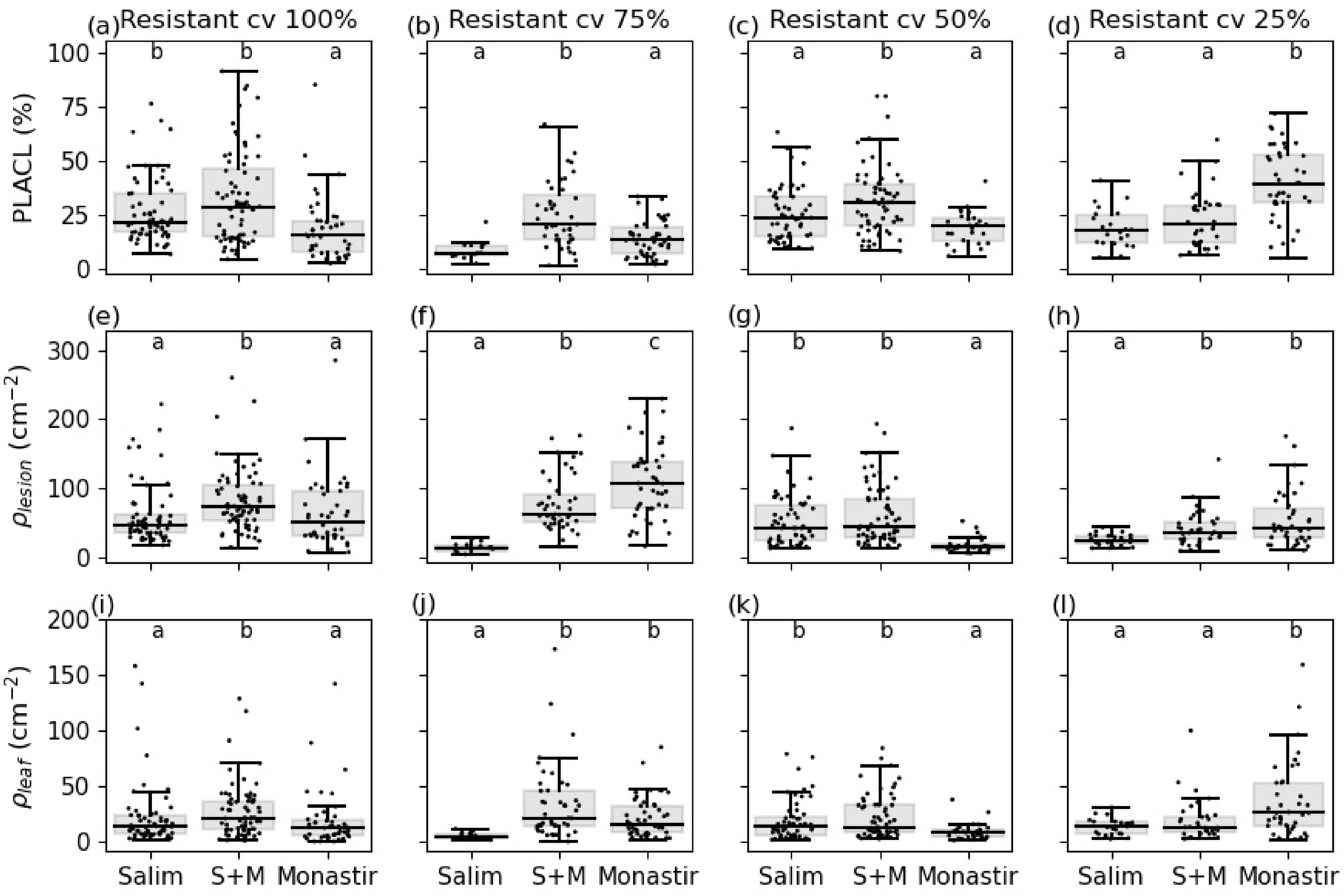


**Figure S15.** The effect of the number of resistant components in the mixture. STB severity at $t_2$ (flag leaves) of the first year. Combinations of the resistant cultivars in equal proportions do not reduce STB severity compared to mixtures with only one of the resistant cultivars.

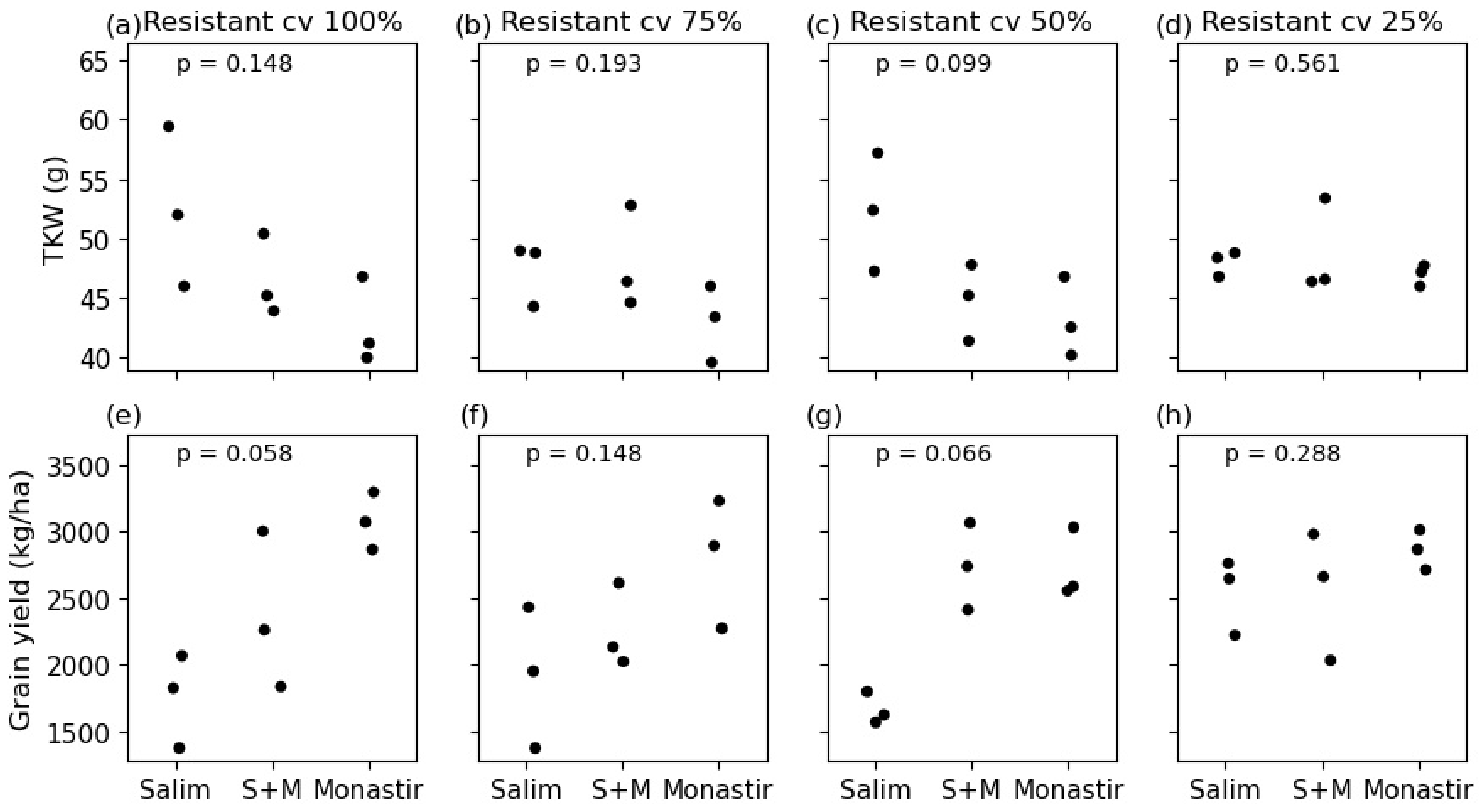


**Figure S16.** The effect of the number of resistant components in the mixture on yield in the first year. Combinations of resistant cultivars in equal proportions do not increase yields compared to mixtures with only one of the resistant cultivars. TKW (a-d), grain yield (e-h). Global Kruskal-Wallis test indicated no significant differences in any of the panels.

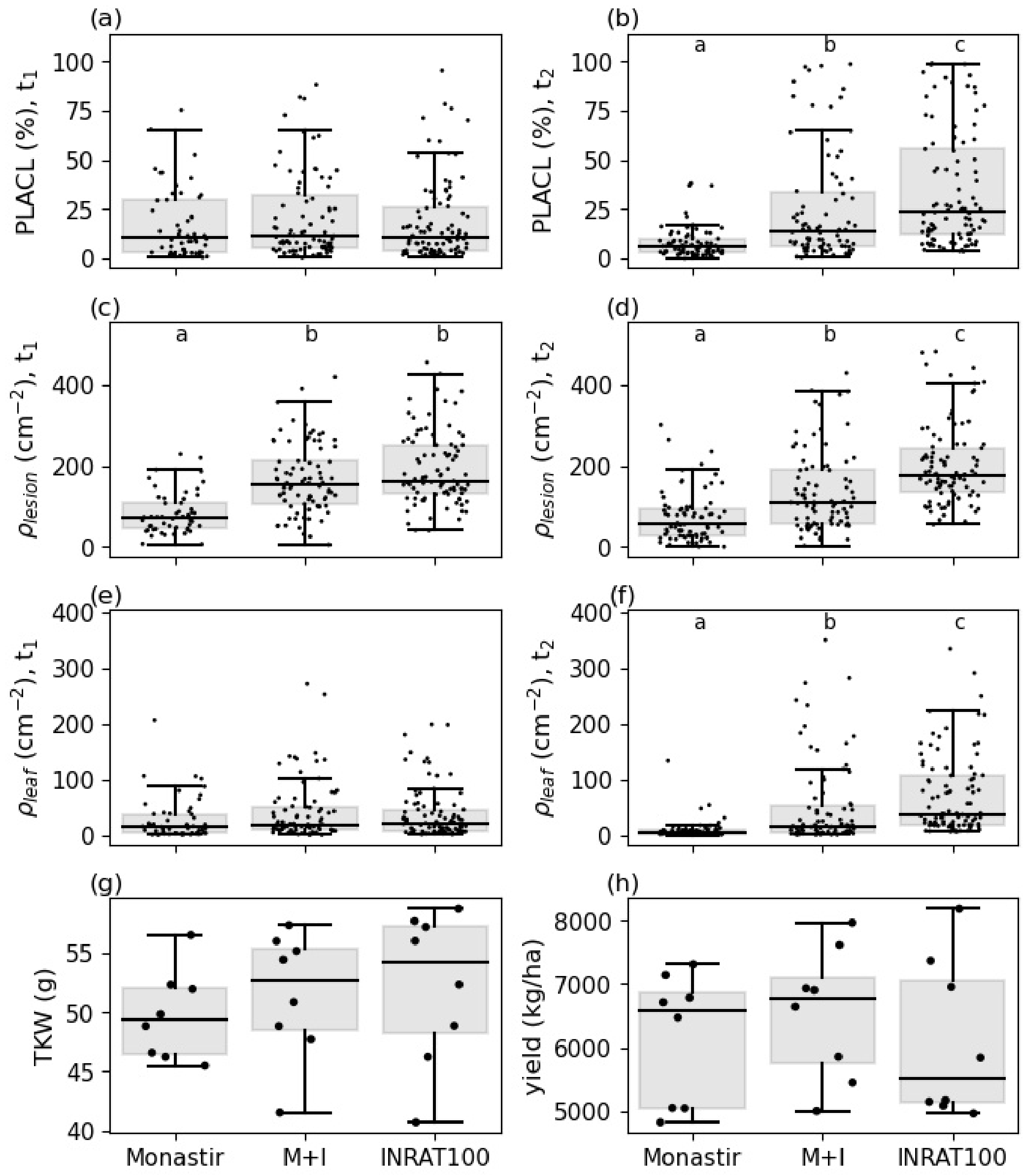


**Figure S17.** The effect of the number of resistant components in the mixture on STB severity and yield in the second year. 1:1-mixture of two resistant cultivars (M+I) do not decrease STB severity nor increase yields compared to pure stands of the resistant cultivars. No significant differences were detected in panels (a), (e), (g) or (h).

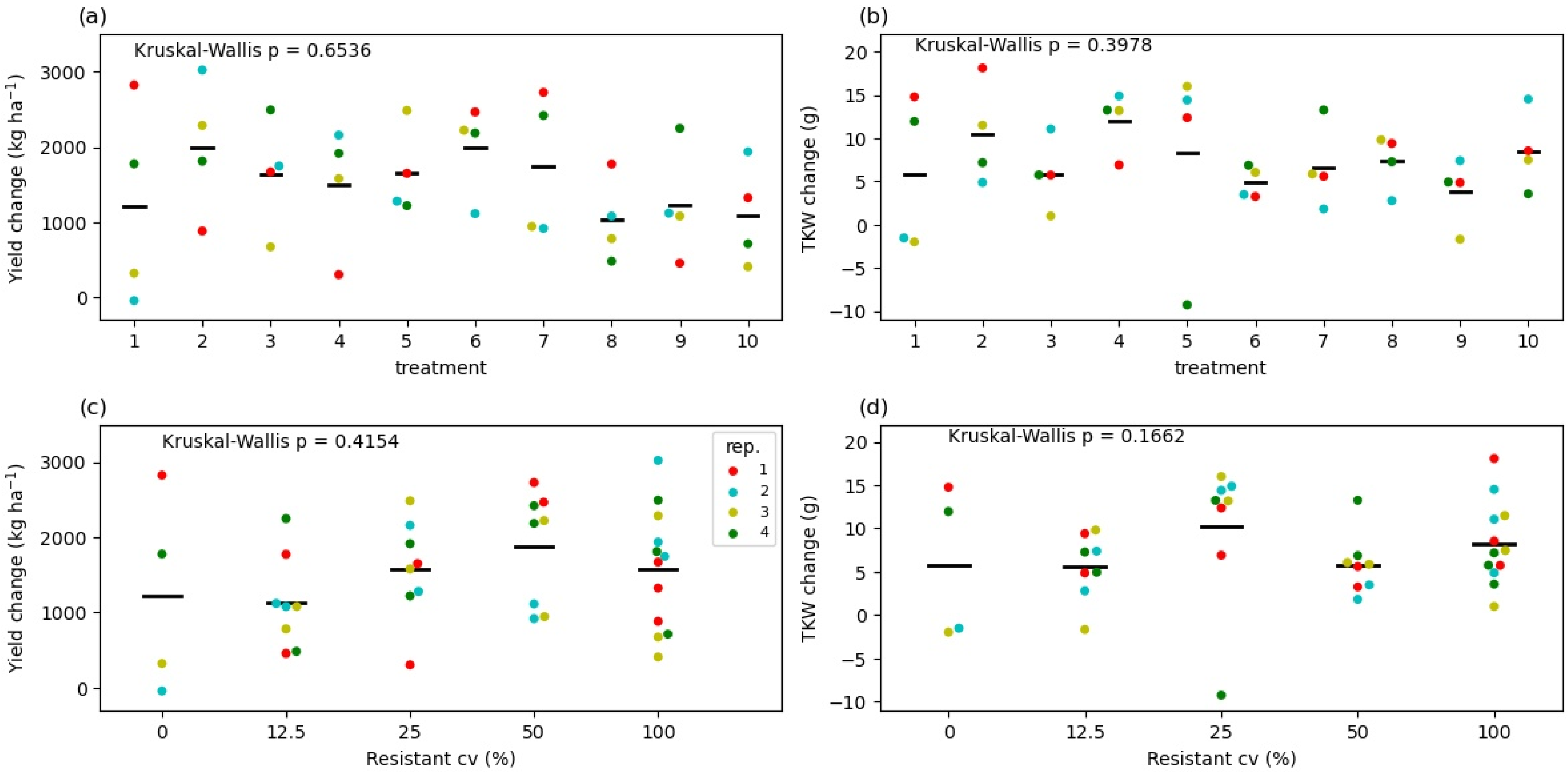


**Figure S18.** Effect of fungicide treatment in grain yield (a, c) and TKW (b, d) in the second year. Each coloured dot shows a comparison between one replicate plot in fungicide treated and one non-fungicide treated plot. Black bar shows mean over the replicates. Bottom row shows data pooled by percentage of the resistant cultivar in different treatment. Global Kruskal-Wallis tests revealed no significant differences between individual treatments nor between resistance levels of the mixtures for either variable.
